# Natural scene segmentation dynamics reveal iterative Bayesian inference

**DOI:** 10.64898/2026.01.30.702842

**Authors:** Tridib K. Biswas, Jonathan Vacher, Sophie Molholm, Pascal Mamassian, Ruben Coen-Cagli

**Affiliations:** Albert Einstein College of Medicine, Dept. of Neuroscience; Université Paris Cité, CNRS, MAP5; Albert Einstein College of Medicine, Dept. of Pediatrics; École Normale Supérieure Paris; Albert Einstein College of Medicine, Dept. of Systems & Computational Biology; Albert Einstein College of Medicine, Dept. of Ophthalmology

**Keywords:** segmentation, psychophysics, Bayesian theories of vision, natural images, perceptual decision-making

## Abstract

The visual system operates by segmenting visual inputs into distinct perceptual objects. Segmentation is dynamic, as revealed by the tempo of perceptual choices and neural activity in visual cortex. Dynamics for natural stimuli however, are poorly understood because natural scene segmentation is ambiguous and subjective. We measured subjective human segmentation maps for natural images using an innovative paradigm, and uncovered richer spatiotemporal dynamics than predicted by current theories of segmentation. To explain these dynamics, we introduced Iterative Bayesian Inference algorithms for segmentation that iteratively integrate visual inputs with the prior expectation for local smoothness. When visual inputs were consistent with such a spatial prior, iterative inference was faster. Such a predicted relationship between spatial prior and inferential dynamics was evident in our data and correctly reflected each individual participant’s spatial biases. We conclude that iterative Bayesian inference sets the tempo for a fundamental function of natural vision.

## Introduction

Segmentation is the task of grouping visual inputs into the objects and parts that compose a scene, and separating those objects from each other ^1^. Performing segmentation supports varied functions of human vision: from reading these words, to navigating the environment, to understanding the relations between objects. In addition, segmentation influences basic processes in visual perception, and may be disrupted in patients with neurologic or psychiatric disorders ^2^. Therefore, identifying the computational principles of human segmentation is crucial for understanding human visual perception and its neural substrates.

Although segmentation feels effortless and instantaneous in everyday vision, signature dynamics have been found in classical grouping experiments that test whether two locations cued on a display belong to the same curve ^3–7^, the same letter ^8^, or the same object ^9–11^. In those experiments, analysis of reaction times (RTs) shows that the larger the distance between two parts of the visual input, the more time it takes to make perceptual grouping decisions ^3–7,9–11^. An influential theory of segmentation posits that this positive correlation between RT and distance arises from how visual attention modulates the activity of retinotopically-organized neurons in the visual cortex. Namely, when attention is directed to a point in the visual field, neurons near the attended–to object in the retinotopic map should become progressively more active ^6,12^ due to the sequential spreading of attention across those neurons. This theory, and related computational models ^6,11,13–15^, are further motivated by recurrent neural dynamics in early visual cortex ^16–18^.

It is unclear however, if current theories can explain the spatiotemporal dynamics of segmentation of natural images, for which segments cannot be unambiguously defined. To address this challenge, we adopted a recent experimental paradigm ^19^ derived from a two–choice “same segment/different segments” task (Fig. 1). This enabled us to measure subjective spatial segmentation maps and thus relate RTs to perceived segments of natural images. Our experiments revealed richer spatiotemporal dynamics than previously observed. RT increased at larger distances for parts of the image perceived in the same segment (a positive RT-distance correlation), as in past reports. But, surprisingly, RT decreased at larger distances for parts of the image perceived in different segments (a negative RT-distance correlation).

**Figure 1.**
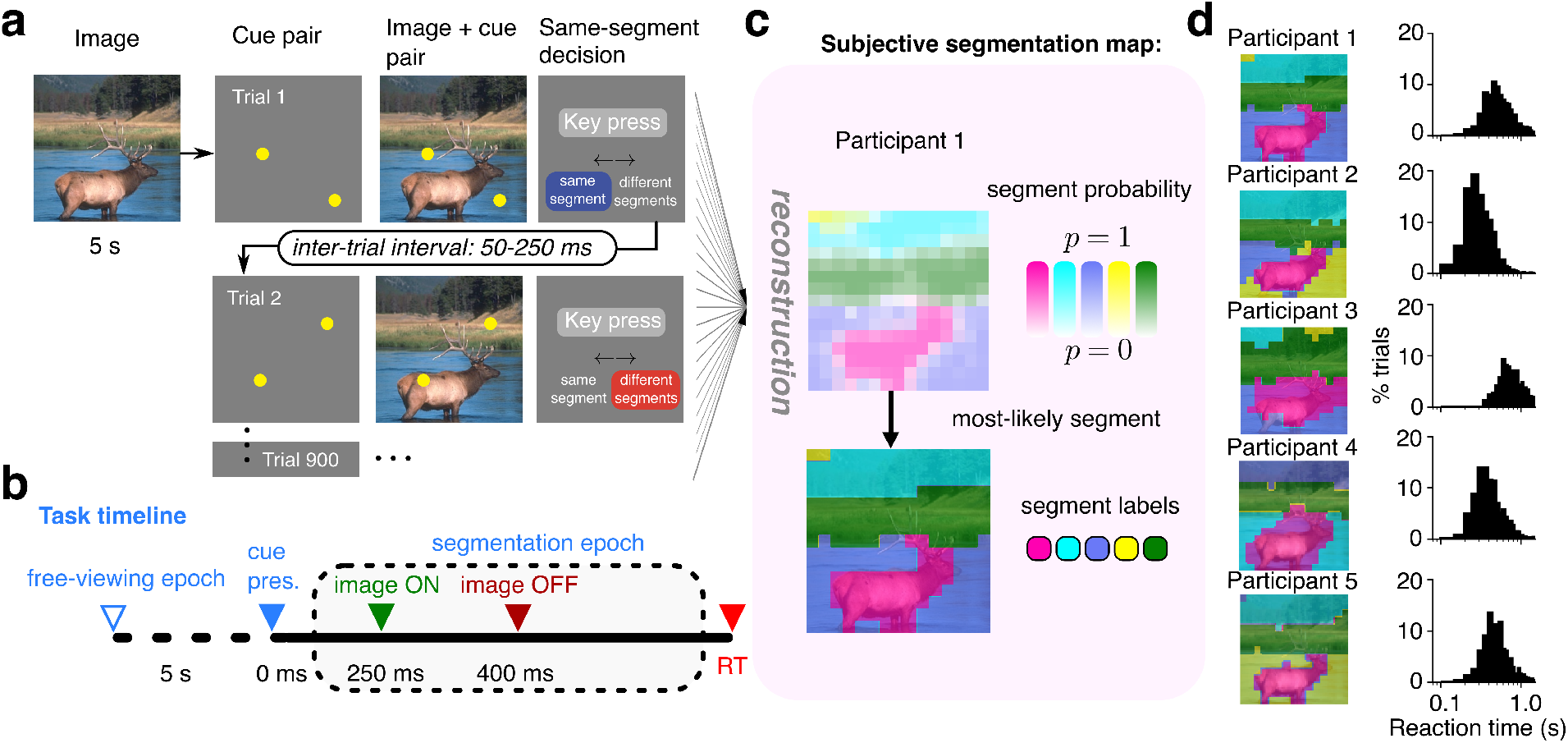
Experimental measurements of subjective segmentation of natural images. **a**, Illustration of one block of the experiment, as described in the main text. During the initial presentation of the image, the participant was asked to mentally partition the image into a pre–defined number of segments (*K* = 5 in this example). Then, in each trial, the participant reported whether the two cued locations were perceived in the “same segment” or “different segments”. **b**, Detailed task timeline. The initial viewing time is designated the “free–viewing epoch”. The time from image onset to response is designated the “segmentation” epoch (illustrated for a single trial). Triangular markers indicate task events. Reaction time (RT) is measured from image offset because participants were not allowed to respond while the image was on screen. **c**, A probabilistic segmentation map (top) is reconstructed from participant responses over many trials (see Methods). Each pixel is colored by the most–likely segment and de-saturated according to segment probability. A deterministic segmentation map (bottom) is derived by assigning the most likely segment/color to each pixel. **d**, Left, deterministic segmentation maps are qualitatively similar across participants (colors are arbitrary and should not be compared between maps). Right, reaction time histograms (*N*_*T*_ = 900 trials) for all five participants in **d.** Bin size = 48 ms.

Rather than proposing an ad–hoc model to fit the opposite correlations for each response type, we used a normative approach asking *why* such dynamics would arise. We theorized that segmenting natural images requires coupled inference about image segments and image features ^20–23^. This is because knowledge of the segments of an image helps improve the inference of the features (e.g. each segment is expected to have a particular appearance), and, in turn, knowledge of the features helps infer the segments (e.g. similar features are expected to belong to the same segment). Therefore, inference should proceed iteratively: starting from an approximate segmentation, the inference of features is updated according to those segments, then segments are revised according to newly inferred features, and so on until the quality of inference can no longer improve. This visual inference is probabilistic and requires integrating visual inputs with prior expectations ^24^ about features and segments of natural images. We refer to this theory as Iterative Bayesian Inference (IBI).

As our theoretical analysis demonstrates, IBI theory predicts that when visual inputs are consistent with prior expectations, fewer iterations are required to improve the quality of inference. This prediction correctly captures that participant RTs increase with distance within the same segment (because nearby parts of an image are expected *a priori* to belong to the same segment), but decrease with distance between different segments (because distant parts of an image are expected to belong to different segments). Further, in accordance with IBI, the strength of the spatiotemporal effect reflected participants’ choice biases measured independently of RT. Lastly, we ruled out the alternative that segmentation dynamics could be explained by the process that reads out the inferred segments, rather than by the dynamics of iterative inference.

Our results establish IBI as a fundamental computational principle in perceptual segmentation. Iterative algorithms like IBI motivate functional theories of neural and perceptual dynamics ^25–31^ and are abundant in both classical ^32^ and modern ^13,33^ machine learning. Therefore, our results have broad significance for understanding natural vision.

## Results

### Measuring perceptual segmentation of natural images

We adopted a recent psychophysical method to measure the segmentation of entire images ^19^ (Fig. 1a,b; details in Methods). Participants were instructed to build a mental segmentation map of a natural image, which was displayed for 5 seconds during an initial free-viewing epoch. Then, in each one of several trials (termed segmentation epoch), we presented two spatial cues in random locations on a gray background (250 ms), and subsequently presented these same cues on the natural image (150 ms). After image offset, participants were asked to report with a key press, as quickly and accurately as possible, whether they perceived the two cued regions as belonging to the “same segment” or “different segments”.

For each participant seeing an image, we reconstructed (see Methods) subjective segment assignment probabilities (e.g. Fig. 1c, top; in total *N* = 58 unique participant-image cases across *n* = 21 participants) from the set of responses over many trials. We then generated a segmentation map with a hard assignment of the most likely segment to each pixel. The segmentation maps confirmed that participants perceived meaningful visual objects as distinct segments (e.g. the elk, water, grass, and trees in Figure 1c, bottom). Maps for each image were qualitatively consistent across participants (Fig. 1d, left column and Supp. Fig. 1), but were not identical.

In addition to measuring subjective segmentation maps, we measured per–trial choice RTs. Participant RT distributions were often skewed (e.g. Fig. 1d, right column) as is common in empirical studies and related modeling literature ^34,35^.

These empirical measurements provide a rigorous characterization of the spatial and temporal aspects of subjective perceptual segmentation of natural images.

### Spatial distance and perceived segments influence reaction times

What is the relationship between spatial and temporal aspects of perceptual segmentation? Past work that addressed this question using both synthetic ^3,6,12^ and natural stimuli ^9,11^ found that RTs increased with distances between cues. Unexpectedly, we found only a weak trend between RT and cue–distance across all trials (Fig. 2a, top; 35,013 total trials). In individual cases, this correlation was often not significant (*p >* 0.05 for 27*/*58 cases; Fig. 2b, top, faded gray diamonds). Furthermore, among significant cases, positive and negative correlations were equally common (Fig. 2b, top, black-bordered diamonds). The mean Spearman correlation across all cases was: −0.0024, s.e.m. *±* 0.017.

**Figure 2.**
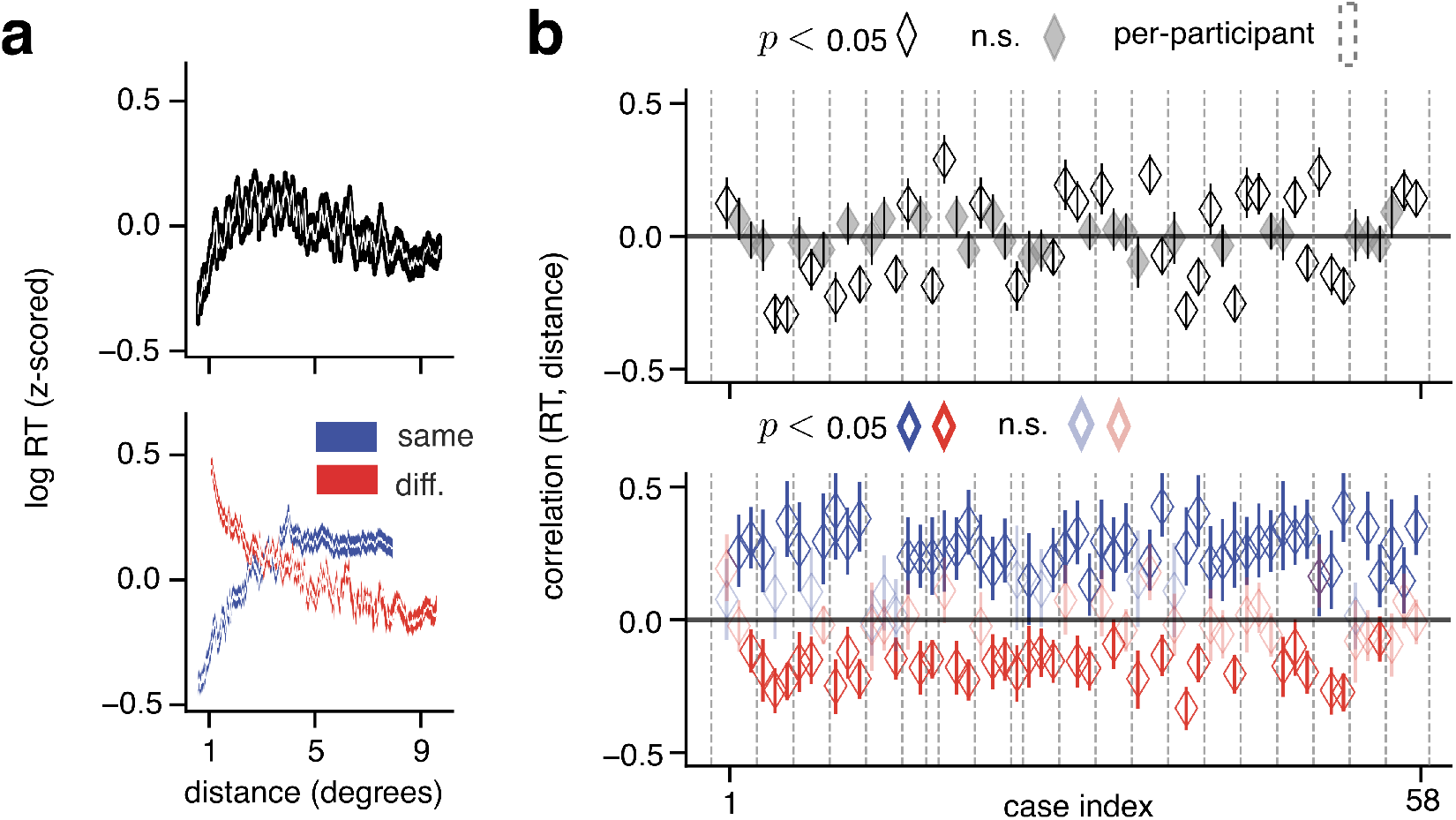
Opposite reaction time-distance correlations within versus across segments. **a**, Aggregate across all cases (one case equals one image and participant). Top, *z*-scored log of RT plotted against distance (in degrees of visual angle) for all trials, sliding window kernel of 600 trials. White lines: mean. Shaded areas: s.e.m. Bottom, trials grouped by whether participants responded “same segment” or “different segments” in a given trial. **b**, Results per case. Top, black, unfilled diamonds indicate the Spearman correlation between RT and distance for cases with significant correlation (*p <* 0.05); gray, filled diamonds for cases with *p >* 0.05. Bottom, blue diamonds indicate the Spearman correlation for the “same segment” response subset, per case, while red diamonds indicate it for “different segments” response subset. Cases with *p >* 0.05 are plotted using faded colors. In both panels, error bars: 95% confidence interval; markers within vertical dashed boundaries indicate cases from a single participant.

This weak RT–distance correlation might appear to contradict past results. Past studies however, focused on cues placed on the same experimentally–defined object. Therefore, we repeated the correlation analysis for only the subset of trials in which participants responded “same segment” (*n* = 10,799 total trials). Consistent with past findings, in this subset the RTs tended to increase with distance (Fig. 2a, bottom, blue line). In all individual cases with significant correlation (*p <* 0.05; *N* = 46*/*58) the correlation was positive (Fig. 2b, bottom, blue diamonds). Across all cases, the mean correlation was: 0.23, s.e.m. *±*0.013.

For the complementary subset of trials in which participants responded “different segments” however (*n* = 24,214 trials), there was a decreasing relationship between RT and distance (Fig. 2a, bottom, red line). In all significant cases (*N* = 34*/*58) the correlation was negative (Fig. 2b, bottom, red diamonds). Across all cases the mean correlation was: *−*0.092, s.e.m. *±*0.014. These findings explain why weak correlations were found when analyzing the complete set of trials. Opposite signs of correlations in the “same segment” and “different segments” subsets canceled out (noting that the “different segments” subset comprised 69% of all trials, but the magnitude of positive correlation was larger for trials in the “same segment” subset).

This result suggests that theories of attentional spreading within an object paint an incomplete picture of the dynamics of segmentation across the visual field.

### A normative theory of segmentation dynamics

To understand these opposite RT–distance correlations, we formulated a normative theory. Such theories assume a central goal of the biological system under study, and ask if algorithms that implement computations toward that goal reproduce the behavior of that system. Here we assumed that segmentation, *i*.*e*. estimating the probability that each pixel is assigned to any possible segment, is a central goal of the visual system. In machine learning, this assignment problem is often solved with a well–established scheme that iterates between estimating the segments and estimating the visual features that comprise each segment, until convergence ^37^. We hypothesized that the visual system uses a similar probabilistic clustering strategy, tuned to the structure of natural scenes, using a Bayesian prior that favors spatial smoothness (*i*.*e*. nearby visual inputs belong to the same segment). We termed this theory IBI.

### Iterative Bayesian Inference explains qualitative spatiotemporal trends

To test this theory, we introduced a model that considers the entire psychophysics task time-line (Fig. 3a,b), and models the segmentation epoch using a segmentation algorithm that implements IBI ^38^ followed by a readout that produces model choices and RTs (Fig. 3c-f, detailed in Methods).

**Figure 3.**
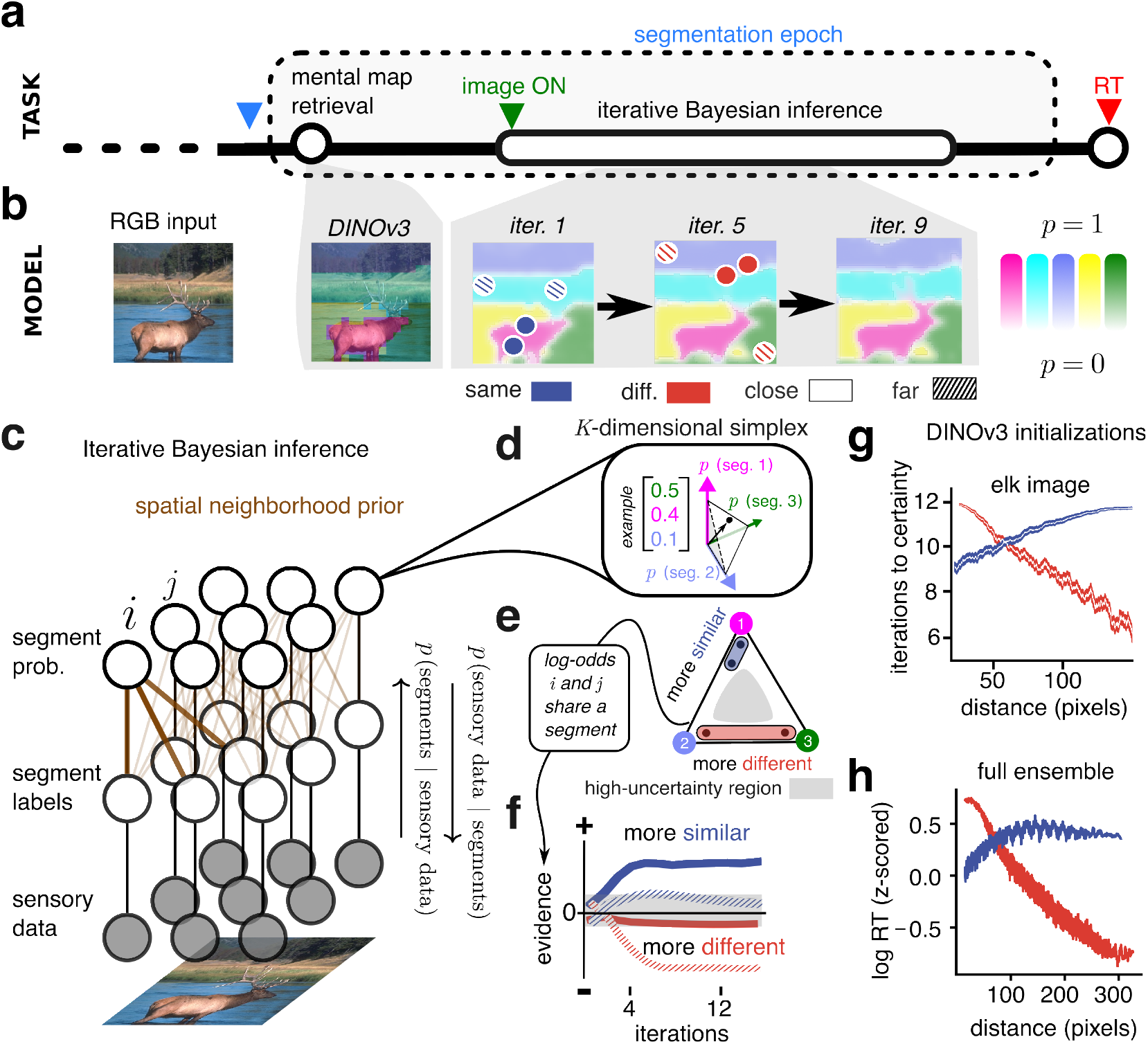
The IBI model qualitatively predicts opposite reaction time–distance correlations. **a**, The timeline of our task. Circular and elliptical markers indicate the temporal position and extent of the processes that the model assumes throughout the segmentation epoch. **b**, Model implementation of each timepoint in the task. The mental map is modeled as the segmentation map generated by DINOv3 ^36^. Prototypical examples of “same” / “different” pairs (blue and red circles) at near and far distances (filled versus hatched) are shown. De-saturated colors in the maps indicate higher uncertainty. Colors are not semantic or ordered, as in Fig. 1b. **c**, In IBI theory, sensory data extracted from the pixels of an image (bottom) is probabilistically assigned to a segment label (middle) with a corresponding probability (top). Inference of the segment assignment uses a probabilistic graphical model, with nodes indicating random variables and edges indicating probabilistic relationships. Brown edges encourage nearby pixels to share the same label. Right: Arrows indicate information passed on the graph during two steps in which labels are inferred from the image features and segment probabilities of the previous step, and then probabilities and features are updated based on labels until convergence (see Methods section “Iterative inference” and Supp. Note 14D). **d**, The segment probability nodes are vectors (each component is the probability that the pixel is assigned to one of *K* segments) that are constrained to the *K*-dimensional simplex. Three dimensions are used in the visualization. The small black arrow and dot indicate an example vector with the probability values listed. **e**, The probability of two pixels being in the same segment can be judged by how close simplex vectors are to each other. **f**, Model readout: Computing the log–odds for two pixels *i* and *j* at each iteration leads to a scalar evidence (ordinate). Example pairs and prototypical evidence for them are shown, following the legend of **b. g**, Number of iterations required to surpass the certainty threshold, plotted against distance, for 25,200 pairs from the image of **a**; sliding window kernel of 800 trials. White lines: mean. Shaded areas: s.e.m. **h**, The *z*-score of the log of the number of iterations required to surpass the certainty threshold plotted against distance, rolling average of 2,116,800 pairs with a sliding window kernel of 800 trials. *z*-scoring is performed per image.

Given an input image, the segmentation algorithm converges to a most–likely configuration of segments starting from an initial guess. We modeled the initial guess as analogous to the participant’s mental map, which is degraded due to visual working memory limitations and must be refined in order to make spatial judgments. Because segmentation of natural images largely reflects meaningful objects, we generated plausible model initializations using a vision transformer (DI-NOv3, ^36^) whose segmentation maps capture object semantics ^11^ (Fig. 3b, second panel; see Supp. Note 3 for implementation details).

From initialization, segment probabilities for each pixel are updated via eqs. 1 and 2 (see Methods). Figure 3c visualizes this process as a graph where nodes represent latent variables (*i*.*e*. segment labels and their probabilities) and edges represent probabilistic dependencies between those variables. Edges that connect neighboring nodes (Fig. 3c, thick brown lines) encourage pixels in the same neighborhood to share a segment label.

The segment probability node of each pixel is a vector constrained within a space called the probability simplex because probabilities must sum to 1 (Fig. 3d). To arrive at a model response in the segmentation task for a pair of pixels, we computed the pairwise probability that both pixels are in the same segment (eq. 3). This corresponds to how much their probability vectors overlap, and how far they are from the center of the simplex where uncertainty about assignment is maximal (Fig. 3e). Lastly, we converted pairwise probabilities to log– odds (eq. 4) to obtain positive or negative evidence for a “same segment” decision ^39^. The per-pixel probability vectors change at each iteration of inference, thus updating the evidence (Fig. 3f). Our model produced an output when the magnitude of evidence exceeded a threshold denoting high certainty. The corresponding number of iterations was the model RT.

Figure 3f illustrates the average dynamics of evidence for the example elk image, while Fig. 3g illustrates how this leads to a spatiotemporal trend. This trend is conserved across an ensemble of images (Fig. 3h; see Supp. Fig. 3 for individual examples). By inspecting example pairs in Figure 3b and their evidence traces in Figure 3f we can see that pixels in a spatially homogeneous segment (e.g. the elk’s body) reached certainty faster than pixels at a spatially heterogeneous segment region (e.g. the elk’s horns). These dynamics illustrate a distinctive property of IBI theory: if two points are close, both the initial guess and the graph connectivity increase the prior probability that they are assigned to the same segment. If those points are in the same object, the corresponding features are likely correlated and so few iterations are needed to reach certainty because the evidence aligns with the prior expectation. Conversely, if those points are in different objects (e.g. across segment boundaries), their features are less correlated and thus provide evidence against the prior, increasing the number of iterations. Similar logic applies to points that are far away, except they are initially less influenced by the spatial prior, but become influenced as segment labels propagate throughout the image.

Therefore, without fitting to data, IBI makes a qualitative prediction that captures our key experimental finding; namely, the dynamics of inference are faster for neighboring pixels that belong to the same segment, and also faster for far–removed pixels that belong to different segments (Fig. 3g,h).

### Iterative Bayesian Inference can be tuned to individualized segmentation dynamics

While the normative theory correctly predicts the opposite RT-distance correlations, those correlations varied substantially across participant-image cases in the data (Fig. 2). We asked if the model could capture this heterogeneity. To facilitate this per–case tuning we coupled our IBI algorithm (now designated the encoding phase; Fig. 4a; orange box) to a decoding phase (green box) in which we fit two interpretable parameters — one for evidence reaching a threshold, and another for evidence having converged (Fig. 4b). To tune the parameters we minimized the difference between RT histograms of the model and of the data, computed separately per response type (details in Methods).

**Figure 4.**
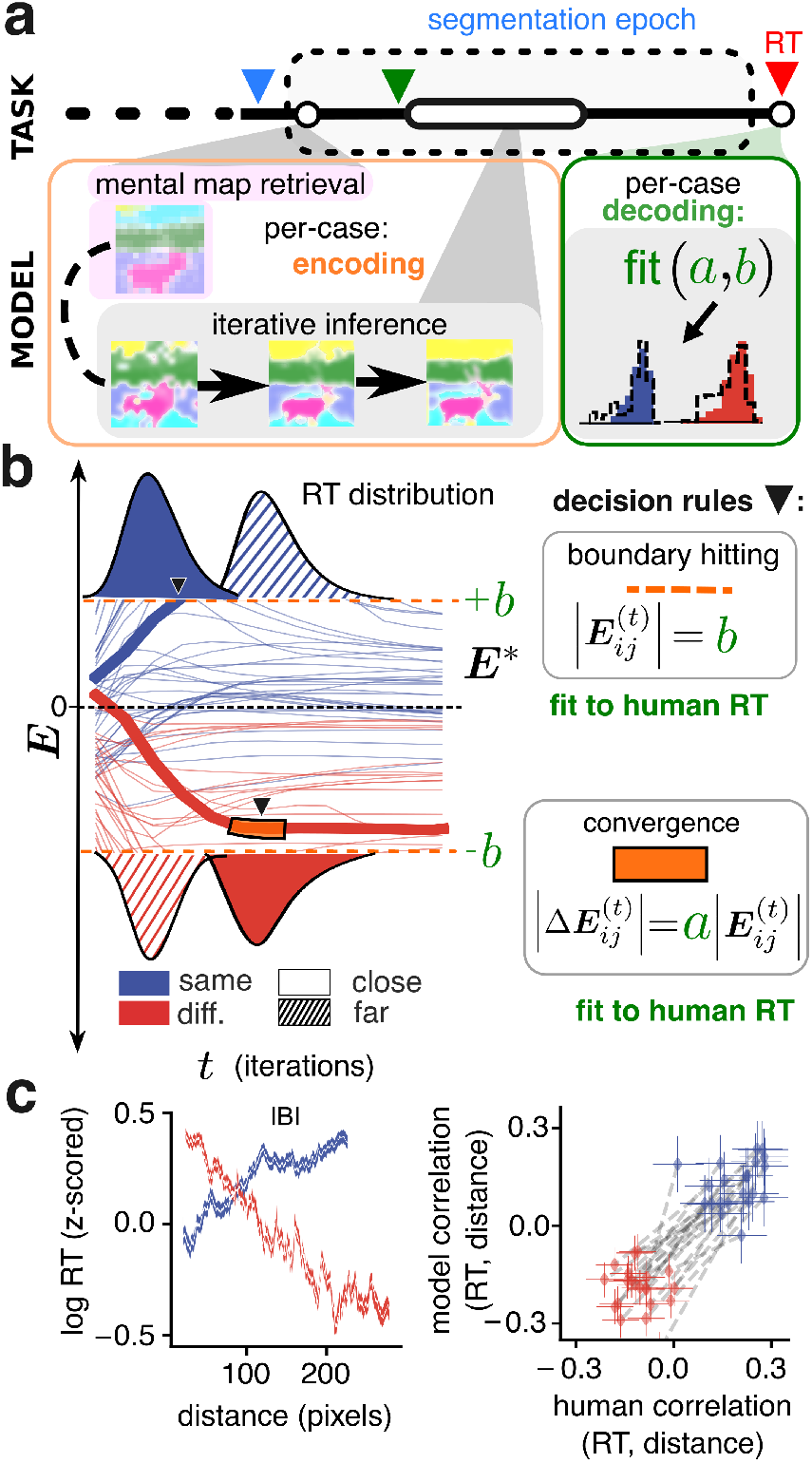
The IBI model accurately fits participants’ RT histograms. **a**, A per– case encoding process (orange border) uses the subjective segmentation map as the initial guess (pink highlight) for iterative inference. A per–case decoding scheme (green border) uses two parameters (*a, b*) to generate model RT histograms (solid dashed lines) matched to the RT histogram for that specific participant (blue filled histogram for same segments decisions; red filled histogram for different segments decision. **b**, Left: Thin blue and red lines illustrate the dynamics of evidence for “same segment” and “different segments” pairs of pixels. The thick lines exemplify how a reaction time is recorded (marked by downward black triangles) when the evidence reaches a threshold value (orange dashed line) or stops increasing (orange highlights, black borders). Top and bottom: mock RT histograms; close pairs = solid, far pairs = hatched. Right: To relate model reaction times to human reaction parameters *a* and *b* are fit (see Methods). **c**, Left: Same conventions as in Fig. 3g, but for the IBI model, aggregated across all cases. Right: Blue diamonds indicate the time-distance correlation for the “same segment” responses subset for each participant across all images they were presented with. Red diamonds indicate the time–distance correlation for the “different segments” response subset. Gray dashed lines connect the diamonds for subsets that belong to the same participant. Error bars: 95% confidence interval.

After fitting, we again examined model RTs and how they trended with distance. Similar to the human data, the model produced opposite RT-distance correlations for the two response types (Fig. 4c). The average correlation across all cases was 0.13, s.e.m. *±*0.017 for the “same segment” subset versus *−*0.2, s.e.m. *±*0.015 for the “different segments” subset. When combining all trials of both types, these trends largely canceled each other out (correlation: *−*0.04, s.e.m. *±*0.005, across cases). Aggregated for each individual participant, the difference in correlations between the “same segment” and “different segments” trial subsets aligned with human values (Fig. 4c, right).

In summary, by including two fittable parameters, our IBI theory can be tuned to reproduce the dynamics per participant and image. This allowed us to quantitatively compare the theory against alternatives, as explained next.

### Ignoring the dynamics of iterative inference disrupts model tuning to human dynamics

We considered an alternative explanation for the observed dynamics: rather than reflecting the time required for the iterative inference of segments, the dynamics may have arisen from the process of reading out the results of that inference. To test this, we assumed that inference could be amortized; in other words, operations needed for IBI could happen quickly whereas time is used to read out the resulting evidence in the presence of noise (Fig. 5a). This is the construction behind drift–diffusion models (DDMs), which are canonically used in decision–making research to model the accumulation of noisy evidence ^40^ and yield skewed RT distributions. Mathematically, the DDM assumes stochastic linear dynamics within each trial ^34^, in contrast to the nonlinear within–trial dynamics that arise from IBI. Conceptually, iterations in IBI are used to improve the quality of the approximate inference from which evidence is derived, whereas in a DDM iterations are used to average out the noise that corrupts the evidence.

**Figure 5.**
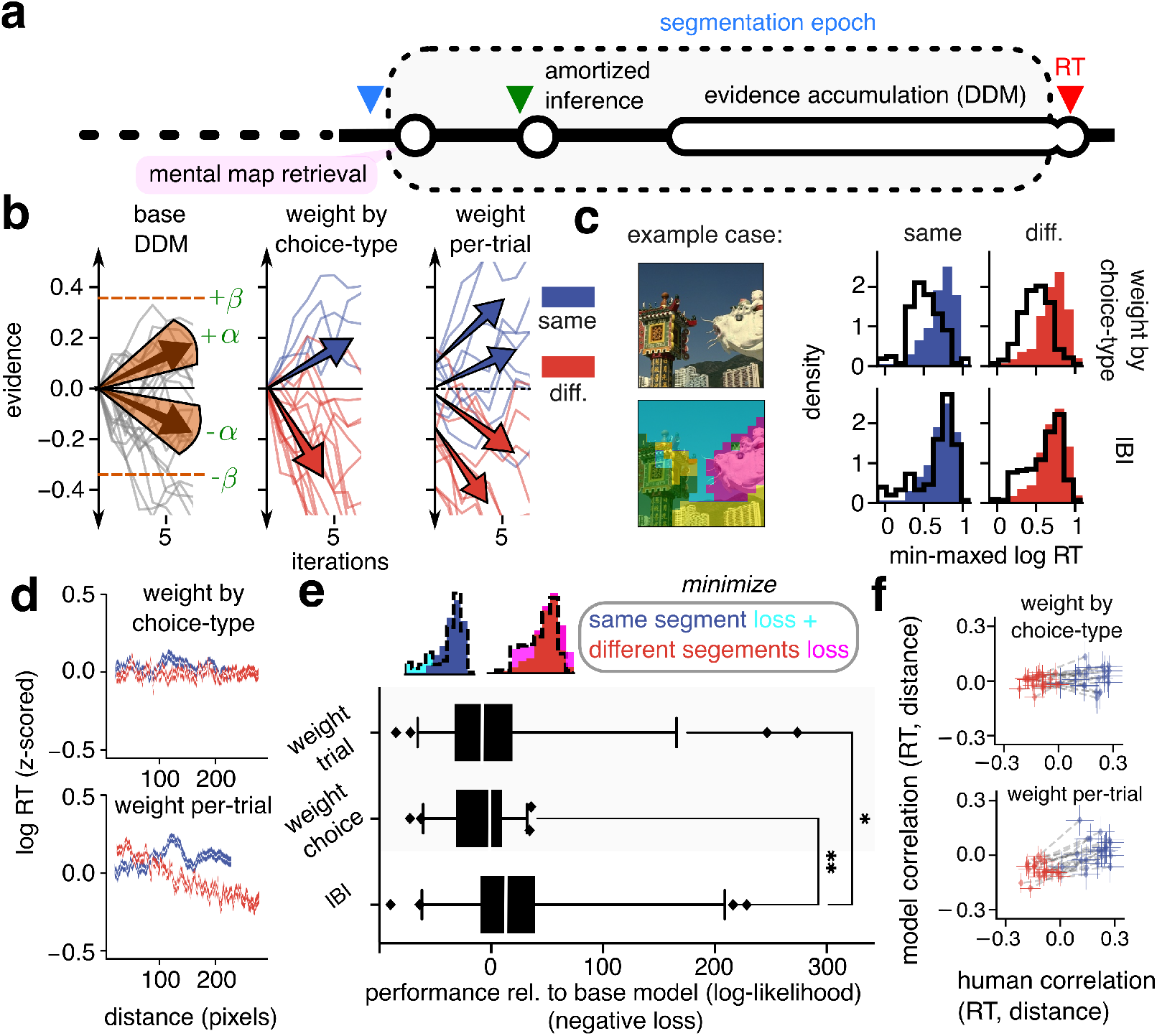
Models that ignore iterative inference do not capture spatiotemporal dynamics. **a**, The alternative models to IBI (i.e. amortized inference followed by evidence accumulation) mapped to task timeline. **b**, Evidence traces from the base DDM (left), a choice-weighted DDM model (middle), and a trial-weighted DDM model (right); DDMs are as described in the main text. Traces are colored by the sign of the trace at the final iteration (not shown) for visual clarity. **c**, Histograms display the fit for the example case from the choice-weighted DDM (top row) vs. the fit from the IBI model (bottom row). Human RT distributions are filled in, while model RT distributions are overlaid as outlines, *n* = 615 trials. **d**, Same as Fig. 4c left, but for the data generated by the model class indicated in the panel title. **e**, (Above) A cartoon depicting the optimization procedure as minimizing histogram mismatch. (Below) Quantitative model performance: Log-likelihoods of all models relative to the base model. Model losses were computed using 5-fold cross-validation. Whiskers indicate 5^th^ and 95^th^ percentile values, the Kruskal-Wallis test with Dunn’s test was used for pairwise comparisons. (*\*p <* 0.03; *\* *p <* 0.003). **f**, Same as Fig. 4c right, but for the data generated by the model class indicated in the panel title.

To establish a reference point, we first constructed a base DDM (Fig. 5b, left) with two parameters, one for the initial value of evidence (starting point) and one for the rate of evidence accumulation (drift rate). As expected, this model did not fit RT data nearly as well as IBI (the IBI performance was on average 13 log– points *i*.*e. e*^13^ times better than base DDM, with cross-validation; Fig. 5e, see Supp. 16 for details). Next, to test the alternative hypothesis and compare IBI and DDM on equal footing, we defined an image-computable, trial–weighted DDM. We used the per–pair log–odds from IBI at convergence to weigh the DDM’s parameters per trial (Methods). Therefore, the trial–weighted DDM effectively included the same information about image feature similarity and the spatial prior as IBI, but ignored the iterations of IBI and instead maintained linear within–trial dynamics. The trial-weighted DDM was not able to fully capture the RT-distance dependence (Fig. 5d and f, bottom) and produced a worse quantitative fit to the empirical data (IBI’s relative cross-validated performance was on average 21.2 log–points better than the trial–weighted DDM; Fig. 5e). We considered also a simpler, choice-weighted DDM, which ignored the IBI evidence but used asymmetric evidence rates (Fig. 5b, middle), to capture that participants in our experiments reacted systematically faster for “different segments” choices (median RT 480 ms, 95% c.i. *±*4.50 ms) than “same segment” choices (512 ms, 95% c.i. *±*5.44 ms; relative difference: 7%; *p <* 1 *×*10^*−*10^). This choice-weighted DDM was on average 15 log–points worse than IBI (Fig. 5e), nor did it improve distribution predictions (Fig. 5c) or distance dependence qualitatively (Fig. 5d, f). Reduced versions of trial–weighted DDMs did not significantly differ from IBI, but they did not capture RT–distance dependence (Supp. Fig. 6). Finally, per case, the IBI model was a better fit than each of the other models in most cases (Supp. Fig. 7).

**Figure 6.**
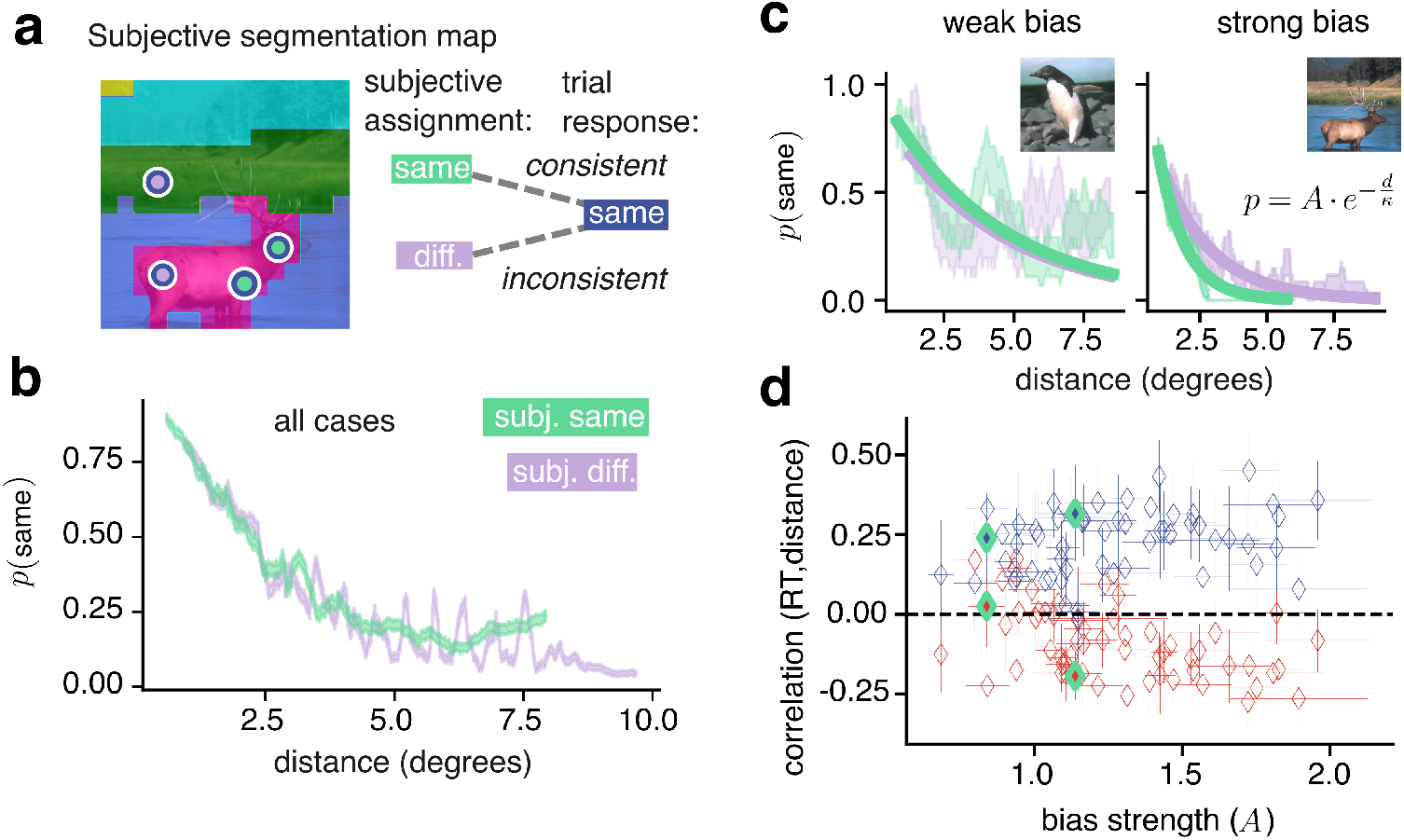
Human response biases reflect a Bayesian spatial prior. **a**, A participant’s deterministic, subjective segmentation map. Markers indicate pairs of cues presented in two trials, both with “same segment” response (blue borders). The response is consistent with the subjective map in one trial (green markers) and inconsistent in the other trial (lavender markers). **b**, Aggregate across all cases and trials, the empirical probability of responding “same segment” as a function of distance (*i*.*e*. the proportion of trials with response “same segment” in a sliding window of 600 trials). **c** Same as panel **b** but for two unique cases, sliding windows of 20 trials (left) and 40 trials (right). Solid lines: Exponential fit to the empirical probabilities, according to the equation in the inset. **d**, Diamonds indicate RT-distance correlation for “same segment” or “different segments” subsets of trials, per case, plotted against the strength of the spatial bias in each case (*i*.*e*. the *A* best fit parameter). The left and right pairs of green-bordered diamonds indicate the left and right example cases from panel **c.** Error bars indicate 95% confidence intervals.

**Figure 7.**
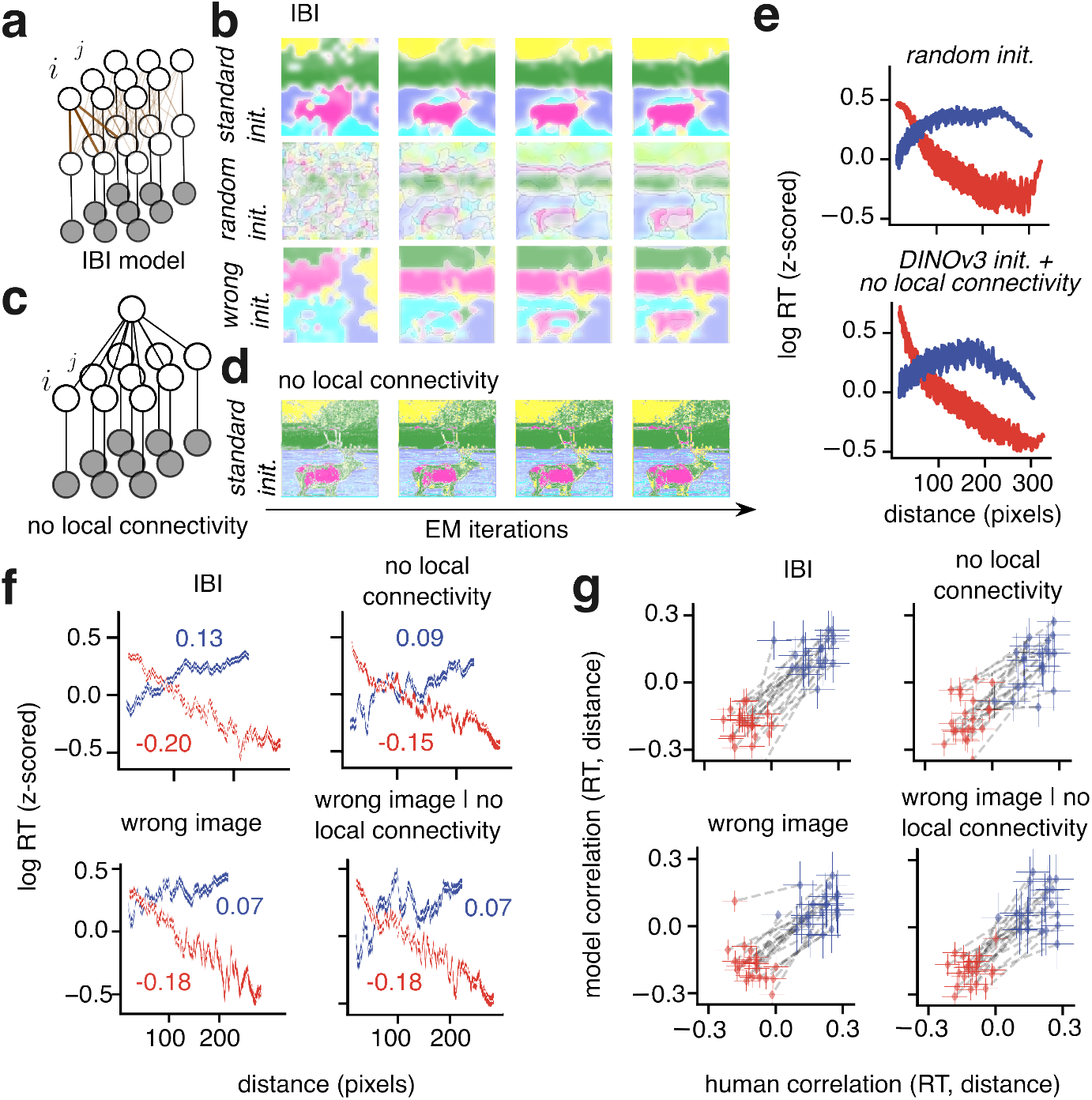
Two mechanisms for the Bayesian spatial prior modulate perceptual segmentation. **a**, The full graphical model (IBI, same as Fig. 3c). **b**, (Top) Iterations in the IBI model with initialization to the participant’s mental map. (Middle) Iterations in the IBI model with a uniformly random initialization of segments. (Bottom) Iterations in the IBI model with initialization to a mental map from the wrong image. **c**, Perturbing local connectivity by replacing the top-layer of segment probability nodes with a single node, *i*.*e*. there is an identical prior probability of segments, for all pixels. **d**, Initial iterations of the model without local connectivity, but with the same initial guess as in panel **b**, top (maps with the wrong image initial guess appear qualitatively the same). **e**,(Top) The log of the number of model iterations needed to surpass the certainty threshold, *z*-scored per image, plotted against distance, for the natural image ensemble of Fig. 3h, initialized with uniformly random segments. (Bottom) The same quantities plotted as above, but the model was initialized with DINOv3 while local connectivity was perturbed **f**, Panels use the same plotting conventions as Fig. 4c, left. The numbers in each panel indicate the Spearman correlation between RT and distance for “same segment” pairs (blue text) or “different segments” pairs (red text) **g**, Panels use the same plotting conventions as in Fig. 4c, right.

From these comparisons, we conclude that the non-linear within–trial dynamics of IBI allow the model to capture aspects of perceptual segmentation dynamics that are not captured by the linear accumulation of noisy evidence.

### Subjective segmentation maps reveal a human spatial proximity bias that reflects a Bayesian prior for segmentation

As explained above, the nonlinear dynamics of IBI are influenced by the Bayesian spatial prior. However, it is possible that human participants did not rely on a spatial prior for their inference of segments, and that instead the RT–distance correlation in the data reflects other unmodeled factors. To address this concern, we required a model–agnostic, experimental measurement of the spatial prior that was independent of the RT patterns. Specifically, we reasoned that if participants relied on a spatial prior, their choices should display characteristic proximity biases.

A proximity bias for perceptual grouping of simple parametric stimuli is well established ^1^. In natural scenes however, it has not been measured before. This is because the subjectivity of natural scene segments^19^ means that a proximity bias cannot be simply measured by the experimenter characterizing responses as correct trials or error trials.

Our experimental paradigm offers a solution. We tested for the proximity bias by studying how participants’ single–trial perceptual responses deviated from their reconstructed mental map. In many trials (*n* =22,738/35,013; 65 %), participant responses were consistent with their subjective map (e.g. Fig. 6a, green), but in a substantial minority of trials (*n* =12,275/35,013; 35 %) the response was inconsistent with the map (e.g. Fig. 6a, lavender). We use the term inconsistent to emphasize there are no objective errors. For instance, the lavender-blue circles in Fig. 6a illustrate a trial in which two regions of the image were reported as being in the same segment, despite being labeled in different segments according to the reconstructed subjective map (an inconsistent response).

We found that same–segment choices, whether consistent or not, were most frequent at short distances between cues (Fig. 6b both lines). Across all trials, the correlation between inconsistency rate and distance was *−*0.83, s.e.m *±*0.004 (*p <* 1 *×*10^*−*10^) for same-segment inconsistencies and 0.97, s.e.m *±*0.001 (*p <* 1*×* 10^*−*10^) for different segment inconsistencies. Note that, in the absence of inconsistent responses the green curve in Fig. 6b would remain at 1 at all distances and the lavender curve at 0. Instead, remarkably, the two curves were near identical, implying a strong bias. Furthermore, if inconsistent responses were due purely to occasional attention lapses or motor errors, their proportion would be independent of the spatial separation between cues. Finally, we found that while inconsistent responses presented the same RT–distance trend (Supp. Fig. 11), they were slower overall (consistent responses: 463 *±* 4 ms, inconsistent responses 517 *±* 5 ms; *p <* 1 *×* 10^*−*10^). These results support the existence of the hypothesized spatial proximity bias for perceptual segmentation of natural images.

Although the proximity bias was a robust feature of our data, its magnitude varied substantially across cases. We quantified this by fitting a decaying exponential function to the proportion of “same segment” responses, as illustrated in Figure 6c (*R*^2^ *>* 0.5 for 53/58 and 43/58 cases, respectively for trials with ground–truth “same segment” and “different segments”). Larger values of the amplitude parameter *A* and smaller values of the spatial decay constant *κ* correspond to a stronger proximity bias. We reasoned that if the bias reflects the use of a Bayesian prior, then under IBI theory the strength of the bias should correlate with the magnitude of the prior’s effects on perceptual dynamics. In line with this intuition, when the bias was stronger, the RT–distance correlations had larger magnitude (correlation of bias versus the magnitude of RT–distance correlation differences: 0.57, 95% c.i. *±* 0.17; *p <* 1 *×* 10^*−*5^), consistent with rational use of the Bayesian prior (Fig. 6d and Supp. Fig. 8). Furthermore, because the DDM alternatives failed to capture the RT-distance correlation (5), the performance gap between those models and IBI increased for cases with stronger spatial bias (Supp. Fig. 12).

These results confirm that the proximity bias we measured experimentally reflects a Bayesian spatial prior for segmentation of natural images, and that the prior influences perceptual dynamics as predicted by IBI theory.

### A Bayesian prior for spatial proximity modulates perceptual segmentation through two complementary mechanisms

Having established the importance of the spatial prior in explaining RTs, we set out to determine the contribution of the two computational mechanisms through which the prior operates in our model, namely the initialization of IBI dynamics and the graph connectivity. We compared model performance when we ablated either or both mechanisms.

First, initializing the model to the DINOv3 semantic segments (Fig. 3) or to the participants mental map (Fig. 4), makes nearby pixels more likely than farther pixels to start in the same segment. To remove this component of the prior, we initialized to spatially random segment probabilities (Fig. 7b, middle row). Second, we ablated the graph connectivity (Fig. 7c,d) to remove the enforcement of the prior throughout iterations.

With either ablation alone, the model reproduced the RT-distance correlation pattern qualitatively (Fig. 7e), albeit with a weaker trend for “same segment” pairs at far distances. With both ablations combined, the model did not converge. We reasoned that, for a specific image and pair of pixels, the effect of the initialization reflects both the initial segment probabilities for those pixels (*i*.*e*. the induced spatial prior) and whether those probabilities are compatible with the similarity between features of that specific image. Since both of these are perturbed by the random initialization, we also considered a different ablation that preserved the spatial prior but disrupted the match between segments and features: we initialized to the segmentation map of a different image (“wrong image” initialization; Fig. 7b; bottom row). The model with both wrong image and no local connectivity still reproduced the qualitative RT–distance correlation (Supp. Fig. 3c). Together, these findings indicate that introducing the spatial prior either at initialization or through graph connectivity is sufficient to capture qualitative spatio–temporal dynamics.

These trends were confirmed after fitting the models to human data. The signs of RT-distance correlations were largely conserved but the magnitude of correlations was lower (Fig. 7f) and less aligned with human correlation values (Fig. 7g), with the exception of random initialization where qualitative trends were perturbed after fitting (Supp. Fig. 4). Comparing the quantitative match between model and human RT distributions, i.e. cross-validated fit performance, was not sufficient to distinguish between the full IBI and ablations (Supp. 5).

For further insight into how the ablations changed our model, we also compared the alignment between model and human responses. Because response alignment was not part of the loss function used to fit RT data, it only reflects the segmentation maps produced by IBI. Qualitatively, the maps were noisier in the model with no local connectivity (Fig. 7d), whereas they were less perturbed but emerged more slowly in the model with wrong or random initialization (Fig. 7b). We quantified these differences by measuring the alignment between per-trial model and human responses, and normalizing it to the alignment between the subjective human map and the per-trial human responses (see Methods).

Table 1 shows that the full IBI model aligns its responses better than wrong or random initialization, and substantially better than the no local connectivity models (all *p* values *p <* 1 *×*10^*−*3^).

**Table 1.** Response alignment: Full IBI model vs. ablations. Values indicate Cohen kappa. Parenthetical values indicate the Cohen kappa as a percentage of the alignment of human responses with their subjective, reconstructed map (Cohen kappa 0.19). All confidence intervals, computed via bootstrapping, are on the order of 1 × 10^−4^

| Initialization: | Subjective segmentation map | Wrong image map | Random |
| --- | --- | --- | --- |
| Smoothing |  |  |  |
| Markov smoothing | **0.16 (87%)** | 0.13 (70%) | 0.12 (66%) |
| No Markov smoothing | 0.067 (36%) | 0.064 (35%) | N/A |

In summary, each one of the two mechanism to enforce the prior is sufficient to capture the RT distribution and RT-distance correlations, although less well than the full model. Only local connectivity is necessary for alignment between the model and participants responses.

## Discussion

Current understanding of perceptual segmentation and its neural basis ^12^ relies strongly on the classical observation that the larger the distance between two locations in an image, the longer it takes to make sameness judgments ^3–5,7,9,10^. We have shown that in natural image segmentation, this classical trend is often inverted when the two locations belong to different segments (Fig. 2). We have proposed a theory that predicts both the classical effect and its inverse (Fig. 3), and explains the variability observed across individuals (Fig. 6). In our theory, dynamics emerge from iteration between inferring the features of the visual input and inferring the segments that those features belong to, with a Bayesian prior that favors local smoothness. Ignoring these structured dynamics of sensory inference, as is common in traditional models of perceptual decision–making, reduces the ability to capture empirical observations (Figs. 5). Our results indicate that iterative Bayesian inference may be a fundamental computational principle for organizing natural visual inputs dynamically.

To reconcile our findings with classical experiments on segmentation dynamics, we highlight differences in experimental design and in the underlying theoretical framework. The first experiment to report increasing RTs with distance used a curve tracing task ^3^. In that study, distance was defined by the length of curve between the two cues, and therefore distance could not be defined for cues on different curves. In one study that modified the task to define distance both within and across segments, the trends that emerged were similar to our findings ^4^. However, motivated by the within-object focus of the sequential attention spreading hypothesis, major subsequent studies emphasized dynamics within the same curve ^5–7^ or within the same object ^9,10^.

We show that the signature finding attributed to attentional spreading, *i*.*e*. a positive correlation between RT and distance when responses are “same segment”, is compatible with our normative assumptions about dynamic, probabilistic segmentation. Because our IBI theory assumes a spatial prior that acts across the entire visual field in parallel (through initialization and recurrent smoothing) we were also motivated to explore “different segment” responses. For these, IBI explains the empirical finding of a negative correlation between RT and distance that cannot be attributed to attention spreading. Given that our theory remains agnostic to how spatially-localized attention interacts with the spatial prior, it will be important to study those interactions for natural stimuli.

Furthermore, by measuring subjective segmentation maps perceived by participants instead of experimentally defining segments, we were able to quantify the extent to which this spatial prior leads to a Gestalt-like proximity bias for each individual case. While such a bias was known for simplified and parametric stimuli ^1^, we have provided the first measurements in natural images (Fig. 6).

Several computational models have been developed to account for segmentation dynamics ^6,10,11,13,14^ but their applications to natural images have been limited. The ‘growth–cone’ model ^6^ worked well for outline images but failed for natural images ^10^, likely because it was not image-computable (*i*.*e*. it did not consider the image features). Other image-computable models were either trained only on object outlines ^13^ and thus not applicable to natural images, or used hand–crafted feature detectors that are not effective for natural images ^14^, with one recent exception ^11^. However, all of the aforementioned models produce positive correlations between

RT and distance regardless of choice type, which is not compatible with our data.

Our IBI theory offers a probabilistic account of perceptual decision–making with natural images. According to probabilistic coding paradigms, neural representations of sensory variables are best understood as approximate Bayesian inference ^41^ that is optimized to natural input statistics ^30,42–44^. Our contributions expand the scope of those principles. First, our model is compatible with Bayesian visual–cortical representation models and substantially extends them to build richer representations of natural images: it performs joint inference of features and segments of entire images via recurrent information flow on a retinotopically organized graph. Second, our analysis reveals that the iterative dynamics required to compute joint inferences are the main determinant of the timing of perceptual segmentation decisions.

Our results do not argue against evidence accumulation and related DDMs as fundamental in decision making. In fact, that framework provides a normative basis for how our model makes decisions using the log–odds. However, most DDM formulations are disconnected from the time–course of inferring the evidence itself. We have proposed IBI as a framework to address this shortcoming and demonstrated that IBI models can indeed be used to define image–computable DDMs. Model comparison demonstrated that a DDM’s ability to capture spatiotemporal dynamics of segmentation improves when it includes image information. However, we also found that within–trial inferential dynamics—which emerge naturally from IBI—are necessary to fully account for the data. There are modified formulations of DDMs that capture non–linear within–trial dynamics, for instance with time-dependent decision boundaries or drift ^29,45,46^. To capture our data, those DDMs would need to be parametrized as a function of both choice type and distance between cues. Rather than deriving new ad–hoc parameterizations of DDMs, our modeling demonstrates the advantage of considering how the visual system represents natural images. We thus show that within–trial dynamics emerge normatively from a theory that encompasses sensory inference and decision making.

Within–trial dynamics in our model also lead to predictions for neural activity. Single–neuron dynamics in V1 align well with the positive correlation between decision time and distance ^12^. Our novel finding, that this correlation can be reversed, raises the question of whether a corresponding signature exists in early visual cortex. In addition, correlates of the spatial prior may be evident in large–scale dynamics of spontaneous ^47^ and evoked ^15^ activity, reflecting the two computational mechanisms needed to integrate the spatial prior in IBI (Fig. 7).

Although we have focused on segmentation, the insights from IBI theory are likely relevant for modeling perceptual and neural dynamics with natural stimuli more broadly given that ecological priors abound in perception ^48,49^, and that natural stimuli often engage non– trivial sensory cortical dynamics ^47,50^.

## Methods

### Experimental procedures

#### Participants

We recruited 21 participants (11 female; age range 16-30 years) with normal or corrected to normal vision. Participants were naive to the task, and they (or their guardians, for minors) gave signed consent to participate in the experiments, and were compensated according to institutional guidelines. The study was approved by the Internal Review Board of Albert Einstein College of Medicine and Montefiore Medical Center (IRB number: 2019–10297).

#### Task details

We used the experimental paradigm and task design validated in our earlier work ^19^. We first provided verbal instructions followed by a brief training session (25 trials). Each participant then completed three “blocks” (we term all trials for one image a block) separated by short breaks. At the beginning of each block, participants were instructed to mentally segment the upcoming image with a predetermined number of segments, *K*, ranging from 3 to 5 (blocks were ordered pseudo– randomly). The image was presented on the screen during a “free–viewing epoch” lasting 5 seconds. Next, a prompt instructed the participant to press a key to start the task. In each trial, we displayed two cues (red circles with 1–degree diameter) on a gray screen for 250 ms and then the same two cues superimposed on the image for 150 ms. Participants were not asked to maintain fixation throughout the trial but eye movements were likely minimal during the brief image presentation time. After the image and cues disappeared, participants were prompted to report whether the two cued locations belonged to the same segment or different segments. The prompt remained on the screen until participants responded with a key press. Participants were instructed to answer as quickly and accurately as possible. The following trial started after a delay of 50-250 ms.

The center of each cue was selected from coordinates defined by a square grid of size 15 *×*15. The grid covered the entire area of the image at lower resolution. Participants’ subjective segmentation maps were decoded at this resolution. For brevity, we refer to each pair of cues with the indices (*i, j*) where each *i* or *j* represents a 2D coordinate. Cues were selected pseudo–randomly with the constraint that the pair was unique in each trial. In each block, we presented the minimum number of trials required to decode the segmentation maps at the desired resolution (a small subset of all possible pairs of cues ^19^) that depends on *K*: *N*_*T*_ = 450, 675, 900 trials for *K* = 3, 4, 5, respectively.

#### Visual stimuli and presentation

We selected 84 natural images manually from the Berkeley Segmentation Database ^51^, and cropped them to patches of size 256*×* 256 pixels. All images were prescribed a single value for *K*. All images were used for qualitative simulations that did not require participant data (Figs. 3h,7e) A subset of 12 images was segmented by 4–6 distinct participants (all images and human maps are shown in Supp. 1). Images were presented on a gamma–calibrated LCD monitor. Participants used a chin rest to maintain a viewing distance of either 67 cm or 96 cm and we rescaled image size to span 8.7*×* 8.7 degrees of visual angle. We combined data for the two viewing distances as no systematic differences were observed.

#### Data preprocessing

In each trial we recorded the binary response *R*_*ij*_ and the corresponding RT *t*_*ij*_ (*i*.*e*. the time between image offset and key press). Before analyzing RTs, we excluded trials with *t*_*ij*_ larger than the 90^th^ percentile (those outlier trials may correspond to attentional lapses). We verified that our main results held without this exclusion (Supp. Fig. 9), and when we controlled for sequential across–trial effects (Supp. Fig. 13). We then *z*-scored log transformed RTs separately for each case (*i*.*e*. one image and one participant) so that the RT trends could be meaningfully compared across participants and images.

#### Reconstructing segmentation maps

We reconstructed the subjective segmentation of an image from only the set of responses to all pairs presented in the experimental block as detailed in our previous work ^19^. Specifically, we estimated the probability that any location on a 15 *×*15 grid is assigned to any of the *K* segments, and termed this grid of probabilities a probabilistic segmentation map (e.g. Fig. 1c, top). This was done by numerically minimizing the squared difference between the measured response *R*_*ij*_ and the probability that the pair *i, j* is in the same segment, averaged over all pairs (we also confirmed that this approach did not generate signal from noise, by shuffling participant responses across pairs; Supp. Fig. 2). From the probabilistic segmentation maps, we computed a deterministic segmentation map by labeling each location on the grid with the segment that had highest probability (e.g. Fig. 1c, bottom).

For per–case initialization coupled to the per–case RT histogram decoding, we used the subjective segmentation map as described above, but instead of assigning to the most–likely segments we perturbed the map by sampling (see Supp. 14D.6 for details).

### Iterative Bayesian Inference models

#### Background

Our IBI algorithm was based on flexible probabilistic mixture models termed FlexMM ^38^. We describe here the four main stages of the algorithm (further details are in Supplement 14).

#### Feature extraction

Given an input image, we first extracted features by passing the image through a deep neural network pretrained for object recognition ^52^. The feature space encodes more abstract information than image pixel intensity, such as edge orientation or texture. We used early-to-intermediate convolutional layers of the network because these have shown alignment with neural activity in visual cortex ^53^. Next, we applied Principal Component Analysis to reduce the dimensionality of each layer and obtained a 12-dimensional representation ***x***_*i*_ at each pixel *i* (mathematical details in Supp. 14A).

#### Generative model

FlexMM assumes a generative model in which each observation ***x***_*i*_ depends probabilistically on a set of latent variables (*i*.*e*. variables inferred from observations rather than directly observed), as summarized by the graph of Fig. 3a in which nodes represent variables and edges represent probabilistic dependencies between variables ^32^. In the graph, each gray node corresponds to the image features ***x***_*i*_ for pixel *i* (Fig. 3a, bottom layer). The variable ***c***_*i*_ represents the segment label that the features of pixel *i* belong to (Fig. 3a, middle layer), and is indexed by *k*. The variable ***π***_*i*_ (Fig. 3a, top layer) is a vector of segment probabilities, *i*.*e*. the probability that the features of pixel *i* belongs to any segment label. Given an input image, these probabilities ***π***_*i*_ are what we aim to infer to obtain the probabilistic segmentation map of the image.

Following previous work relating image statistics to visual–cortical activity ^30,42–44^, we assumed that all the features ***x***_*i*_ within a given segment with label *k* follow a Student-t distribution with parameters collectively denoted ***θ***_*k*_ (see Supp. 14B for the specific parametrization). The spatial prior is enforced by introducing additional edges between ***π***_*i*_ and the ***c*** variables in a local neighborhood (Fig. 3a, brown edges) encouraging pixels in the same neighborhood to share the same segment label.

#### Iterative inference with a spatial prior

From the generative model of FlexMM, we were interested in computing the probabilistic segmentation map ***π*** of an input image. Because exact computation of these probabilities is not tractable, FlexMM uses an iterative approximation scheme that extends the popular Expectation-Maximization (EM) algorithm for probabilistic clustering, to account for the spatial prior as follows. EM iterates to find the values of ***π*** (the organization of segments) and ***θ*** (the organization of features within each segment) that increase the likelihood of the observations ***x***. At iteration *t*, the Expectation step uses the previously inferred values of the ***π***^(*t−*1)^ and ***θ***^(*t−*1)^ to update the approximation of the posterior probability for ***c***, denoted ***γ***^(*t*)^. Then, using the current ***γ***^(*t*)^, the Maximization step updates ***π***^(*t*)^ and ***θ***^(*t*)^ to their new best value, while incorporating the spatial prior by considering the current values of ***γ*** in a spatial neighborhood (full derivation in ^38^ and Supp. 14D):

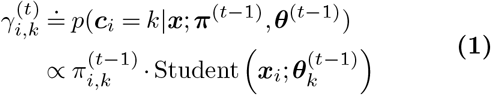

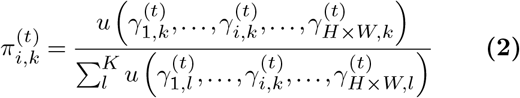

FlexMM implements the spatial prior through the linear function *u* which is a discrete convolution with a 2–dimensional Gaussian kernel of finite size denoted by *H× W* (*H* = *W* in our formulation) centered at pixel *i*. Following from eqs. 1 and 2, the spatial prior was introduced iteratively; the probability of the segment of each pixel was updated by averaging the previously inferred segment probabilities within a spatial neighborhood. Similarly, to further improve segmentation performance, the update of ***π*** at one feature layer took into account also the previously inferred segment probabilities in neighboring layers (see Supp. 14D.5 for details). In addition, we introduced the spatial prior also through the initial guess ***π***^(0)^ for each image and participant, by sampling from the participant’s subjective segmentation map which was often comprised of locally smooth segments (see Supp. 14D.6 for details).

In FlexMM, convergence of the likelihood is guaranteed ^38^ and therefore the dynamics of the probabilistic segmentation map are well defined. EM convergence is typically defined globally for the image (*i*.*e*. on the likelihood summed over all pixels). Yet, the segment probabilities of all pixels do not converge at the same time (mathematical details in Supplement 14D). As explained next, this is important when applying FlexMM to our experimental task.

#### Decision rules for IBI

Because individual pixels displayed unique dynamics, so too did pairs of pixels (see traces in Figs. 3e; 5c,e,h). Our decision rule was based on comparing the probability that a pair of pixels *i, j* are in the same segment (denoted *π*_*ij*_) versus in different segments (1 *−π*_*ij*_). At each iteration, these quantities were computed from the per–pixel probabilities and their log–ratio was used as evidence in favor of one option or the other:

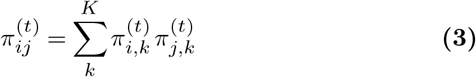

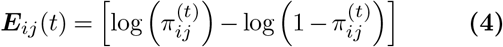

Positive values of ***E***_*ij*_ imply evidence in favor of “same segment” and negative values in favor of “different segments”. The model could report a decision at the first iteration for which the evidence reached a boundary *b* for “same segment” or *−b* for “different segments”. When fitting human data, we treated *b* as a free parameter, and also included a second decision rule, to account for the fact that, for some pairs, *π*_*ij*_ might converge to a value that does not reach the boundary. Specifically, the model could report a decision when the change in evidence between consecutive iterations 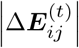 became smaller than the quantity *a* 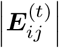, where *a* is the second fittable parameter. We recorded the model decision and corresponding RT at the first iteration in which either rule was satisfied (models that used only a single rule performed worse; Supp. Table 4, Supp. Fig. 10).

### Drift Diffusion Models

#### Decision rules for DDM

Drift rates for the base model were decoded to match participant RT histograms without evidence from the IBI model. For weighted DDMs, drift rates at particular trials were weighted by evidence from the IBI model. For extensive DDM implementation details, see Supplement 15. Because DDMs assume that evidence grows at a constant rate on average, only one stopping rule is needed for when evidence hits the boundary, *β*. We distinguish the boundary for DDM models as *β* because it is independent of the IBI boundary parameter *b*. In addition, we also fit a global gain parameter *α* that scales the evidence for all trials.

### Model fitting (decoding) and comparison

#### Preprocessing

Before model fitting, we normalized RTs per case between 0 (shortest RT) and 1 (longest), separately for the human data and for each model.

#### Parameter fitting

All IBI and DDM models used two free parameters, *a* and *b* or *α* and *β*. Best fit parameters were found by maximizing the likelihood that the observed human RTs come from the distribution of model RTs. We computed such likelihoods separately for the trials in which humans chose “same segment” vs. trials in which the humans chose “different segments”, and used the negative of the sum of those two terms as the cost function to be minimized (equivalent to a maximum-likelihood approach; details in Supp. 16). This cost function was not differentiable with respect to the parameters, so we used Markov-Chain Monte-Carlo (MCMC) global minimization methods (see Supp.16 for details). Note that model parameters were not optimized to capture per–trial RTs or responses. Because our decoder’s ultimate objective is to match histograms, we were able to do so with only two interpretable parameters (see above, Decision rules section).

#### Model comparison

For fair comparison across models we used 5–fold cross validation per case. Models were fit using trials in a training split, and then applied to predict RTs in unseen trials that were part of a test split. All reported results are from applications to test splits *i*.*e*., unseen data.

To compare model–human response alignment, we first used Cohen kappa measure ^54^ to compute the similarity between participant and model responses. We then normalized this Cohen kappa with the Cohen kappa for a participant agreeing with their inferred most–likely response based on the reconstructed map.

#### Implementation details

To add stability to all models we computed the RT for each pair *ij* over 10 pseudo-coordinates that were randomly selected to be within the spatial neighborhood of pixel *i* and *j*. Using 10 pseudo-coordinates provided 100 pseudopairs that ensured small floating-point differences from pixel to pixel did not impact model results.

#### Statistical analysis

To plot the *z*-scored log of RT versus distance, we sorted RTs by distance, and applied a sliding window kernel. When comparing median RTs, we used bootstrapping with 9999 resamples to compute 95% confidence intervals.

We quantified the relation between RT and distance with the Spearman correlation and we used parametric statistics to compute confidence intervals For comparing relative likelihood distributions from each model type, we first confirmed that models were sufficiently different from each other using the non-parametric Kruskal-Wallis test. For pairwise comparisons we used Dunn’s test with a significance level of *α* = 0.05; *p*–values were corrected for multiple comparisons. Comparisons and error estimates for Cohen kappa were via bootstrapping.

To fit exponential curves for quantifying spatial biases (Fig. 6) we used a nonlinear least-squares regression. Parameters *A* and *κ* were optimized to fit a curve with the parametrization *p* = *A·* exp(*− d/κ*), where *p* is the probability of responding “same segment” and *d* is the independent variable (distance between pixels). During fitting, we applied the bounds [0.5, 4.3] (units in degrees) to *κ*. These were chosen as reasonable lower and upper bounds for the space constant given the size of the image. 95% confidence intervals for fit parameters were computed using two standard deviations of optimal parameter estimates.

## Supporting information

segment_probability_panels.mp4

segment_probability_panels_nlc.mp4

segment_probability_panels_rand.mp4

## AUTHOR CONTRIBUTIONS

**T.K.B**.: formal analysis, methodology, software, writing - original draft, writing - review & editing

**J.V**.: methodology, writing - review & editing

**S.M**.: resources, funding acquisition, writing - review & editing

**P.M**.: conceptualization, methodology, funding acquisition, writing - review & editing

**R.C.C**.: conceptualization, funding acquisition, methodology, writing - original draft, writing - review & editing

## ACKNOWLEDGMENTS

The authors would like to thank Carlos Madrid-Aliste for assistance with running FlexMM and parameter fitting on the Einstein High-Performance Computing Cluster. We also thank Adam Kohn, Máté Lengyel, Ralf Haefner, and Odelia Schwartz for productive conversations. For participant recruitment and task administration we thank the study coordinators, Tringa Lecaj and Dennis Cregin. T.K.B. was partially supported by the National Institutes of Health Medical Scientist Training Program (T32-GM149364). R.C.C. was supported by the National Institutes of Health (EY031166; EY030578) and the Rose. F. Kennedy Intellectual and Developmental Disabilities Research Center. R.C.C. and S.M. were supported by the Simons Foundation International (SFI-AN-AR-Pilot-00009855). P.M. was supported by the French Agence Nationale de la Recherche (ANR-19-NEUC-0003-01 and ANR-17-EURE-0017). Support for recruitment of participants was provided by the Human Clinical Phenotyping Core of the NICHD funded Rose. F. Kennedy Intellectual and Developmental Disabilities Research Center (P50 HD105352).

## Supplementary Information

### Supplementary Note 1: Subjective segmentation maps - all images

**Supplementary Figure 1.**
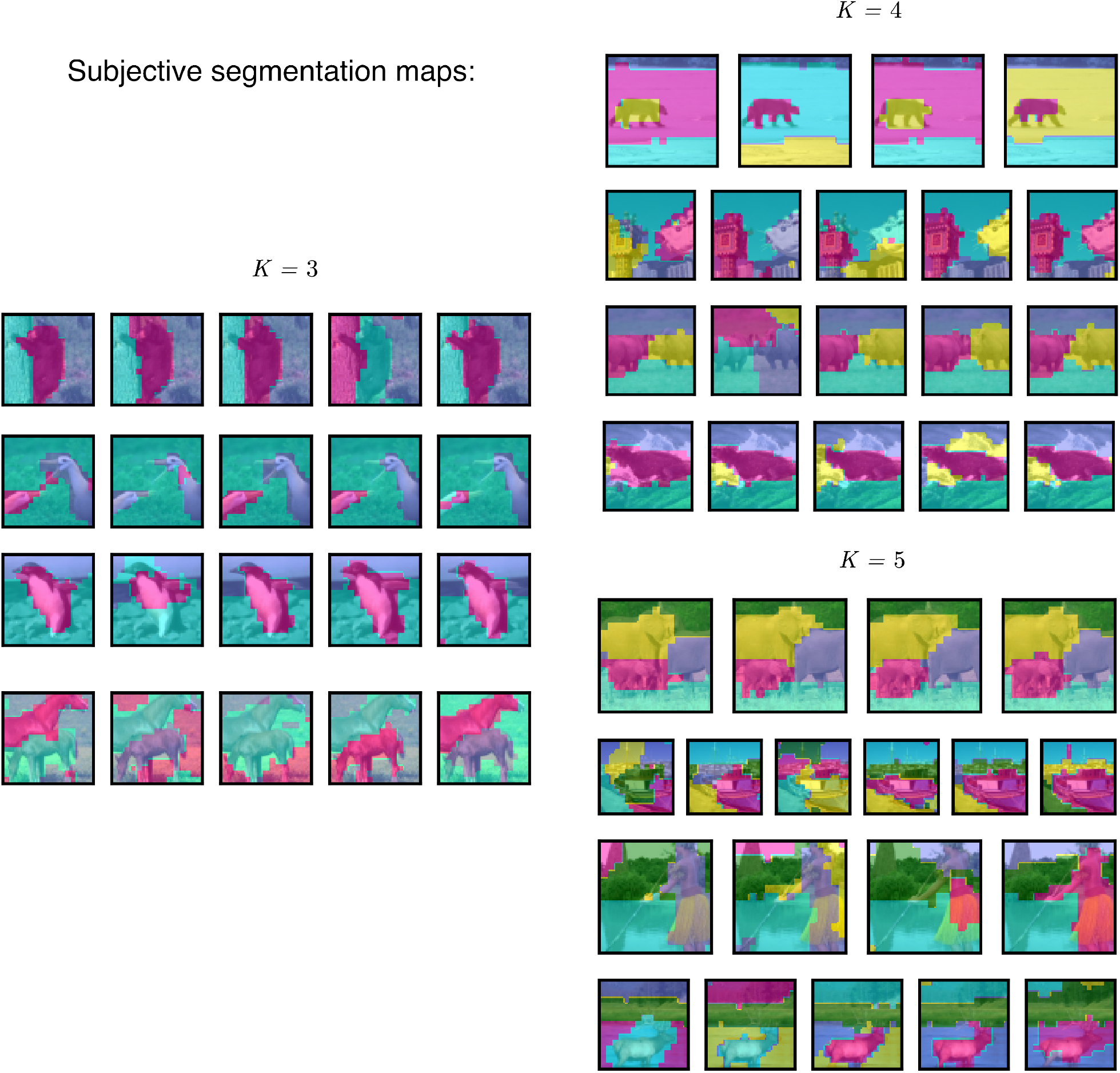
Subjective segmentation maps for all images and participants. Each panel is the segmentation map for one case (one image by one participant). Images are organized by the number of segments requested, *K*. Each region in a 15 *×* 15 grid is assigned the most–likely segment. As in Figure 1b,c.

### Supplementary Note 2: Subjective segmentation map control - all images, randomized responses

**Supplementary Figure 2.**
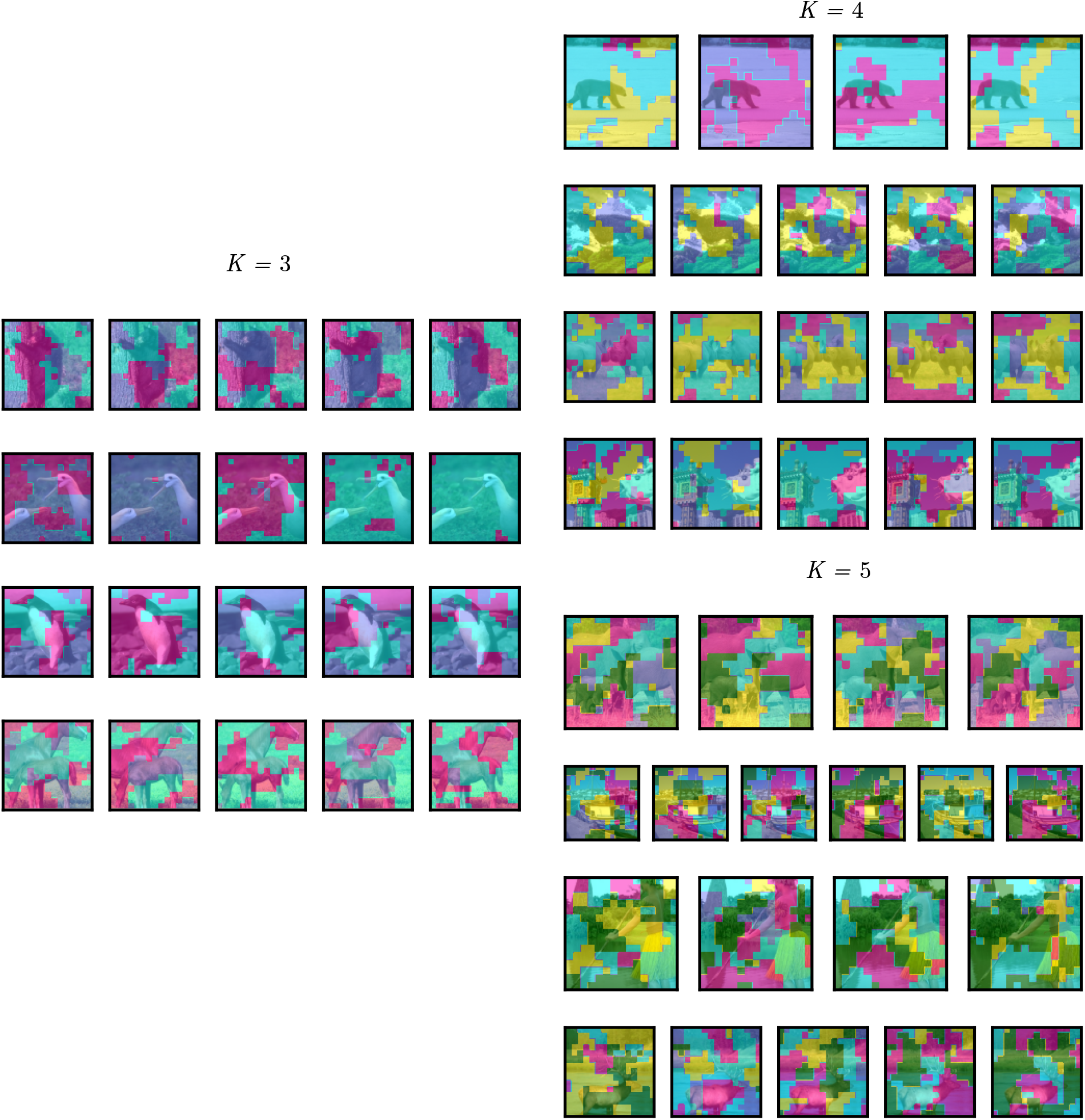
Decoded subjective segmentation maps with participant responses shuffled across trials. We devised a procedure to verify that the decoding algorithm could not hallucinate segments when the binary choice data did not contain segment-related information. To do so, we randomly shuffled the order of the responses across trials, so that each response was not aligned with the actual cue pair shown in the corresponding trial, but it was instead aligned with a different cue pair. This figure shows that, in that case, the maps produced by the decoder were not meaningfully related to the image contents.

### Supplementary Note 3: Qualitative simulations

**Supplementary Figure 3.**
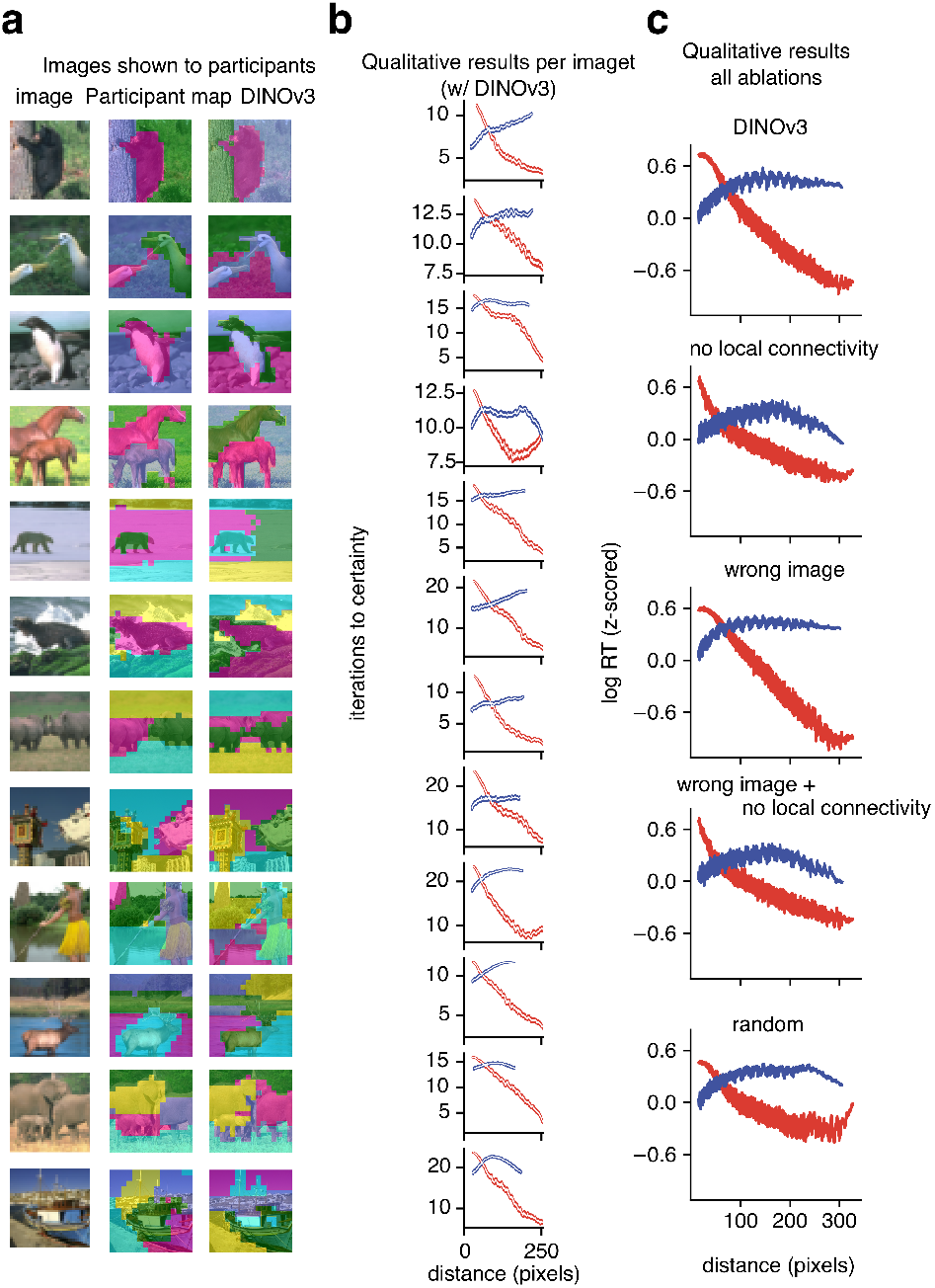
Qualitative simulations with DINOv3 initialization. **a**, Original 12 images presented to participants shown. To create segment labels analogous to a “mental map” without using any information recorded in the psychophysics task we used the pretrained vision transformer DINOv3 ^1^. The patching process of DINOv3 led to maps of 16 *×* 16 grid units, referred to as patches. This was comparable to the human subjective task maps which were 15 *×* 15 grid units. We used the patch features from the last layer of the DINOv3 features, computed the top 6 principal components, and grouped them using a Student-T mixture model (equivalent to FlexMM without local connectivity). **b**, Same plotting conventions as Figure 3g. Using one threshold for certainty across images leads to heterogeneous results across images. **c**, Aggregated across 84 images we see human–like spatiotemporal trends for DINOv3 initialization and various ablations (random initialization, no local connectivity, and wrong image initialization).

### Supplementary Note 4: Random initializations for IBI

**Supplementary Figure 4.**
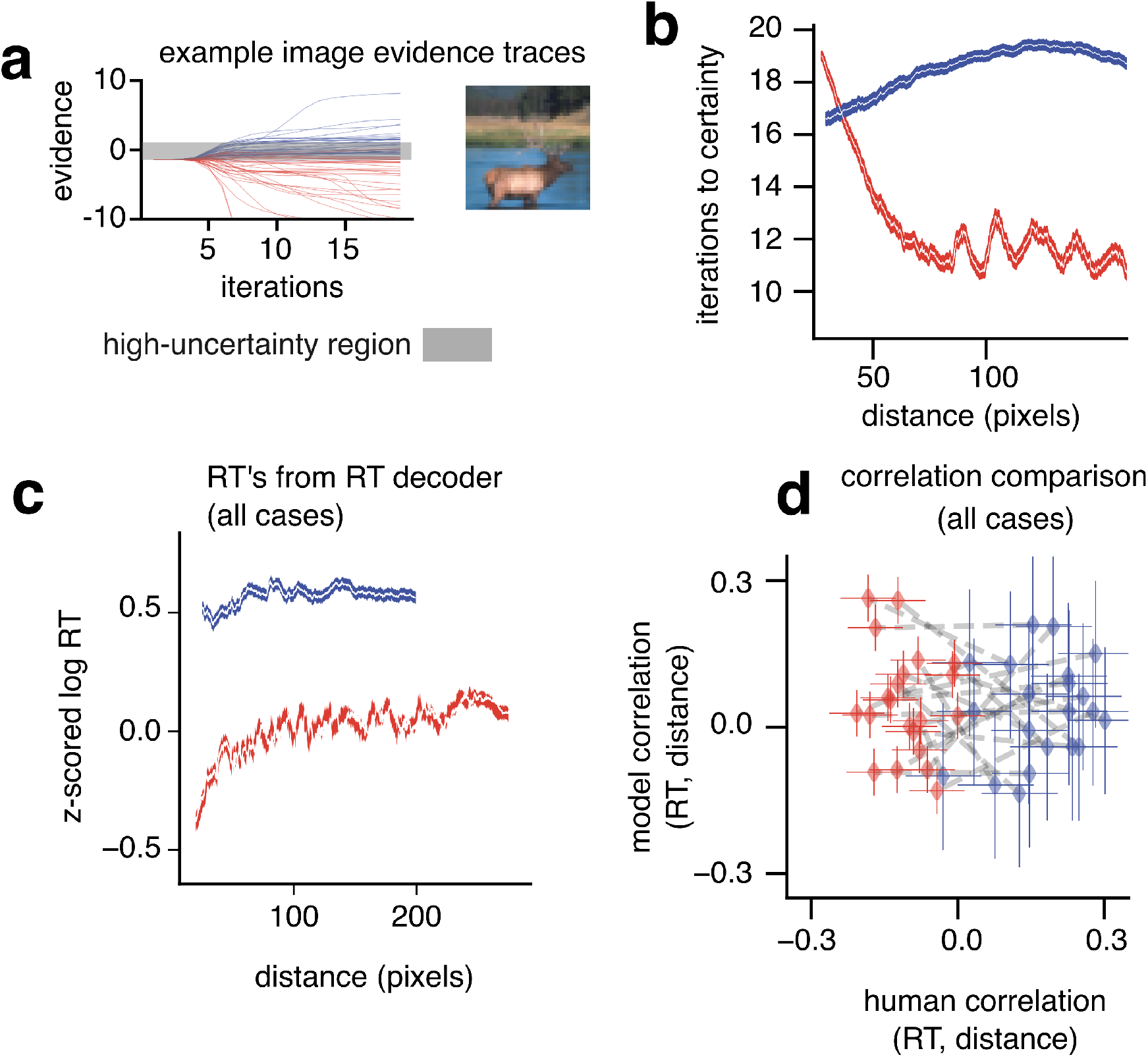
Encoding vs. decoding for a random initialization. **a**, Evidence traces, following plotting conventions in Fig. 3f. **b**, Qualitative results when plotting the number of iterations for the segmentation encoder to reach certainty, plotting conventions follow Figure 3g. Both panels **a** and **b** display results for the example image shown in **a. c**, Plotting conventions follow Fig. 4c (left). **d**, Plotting conventions follow Fig. 4c (right). Both panels **c** and **d** display results for all cases (images and participants) .

### Supplementary Note 5: Model fits - ablations

In the main text, we compared models by subtracting the loss for the base DDM from the loss of other models per each image and participant. This was because we wanted to isolate how much the spatial prior and iterative inference specifically improved quantitative performance, and this could vary across images. In this section, we report absolute values for model fits using our custom loss function (see Supp. 16)

**Supplementary Figure 5.**
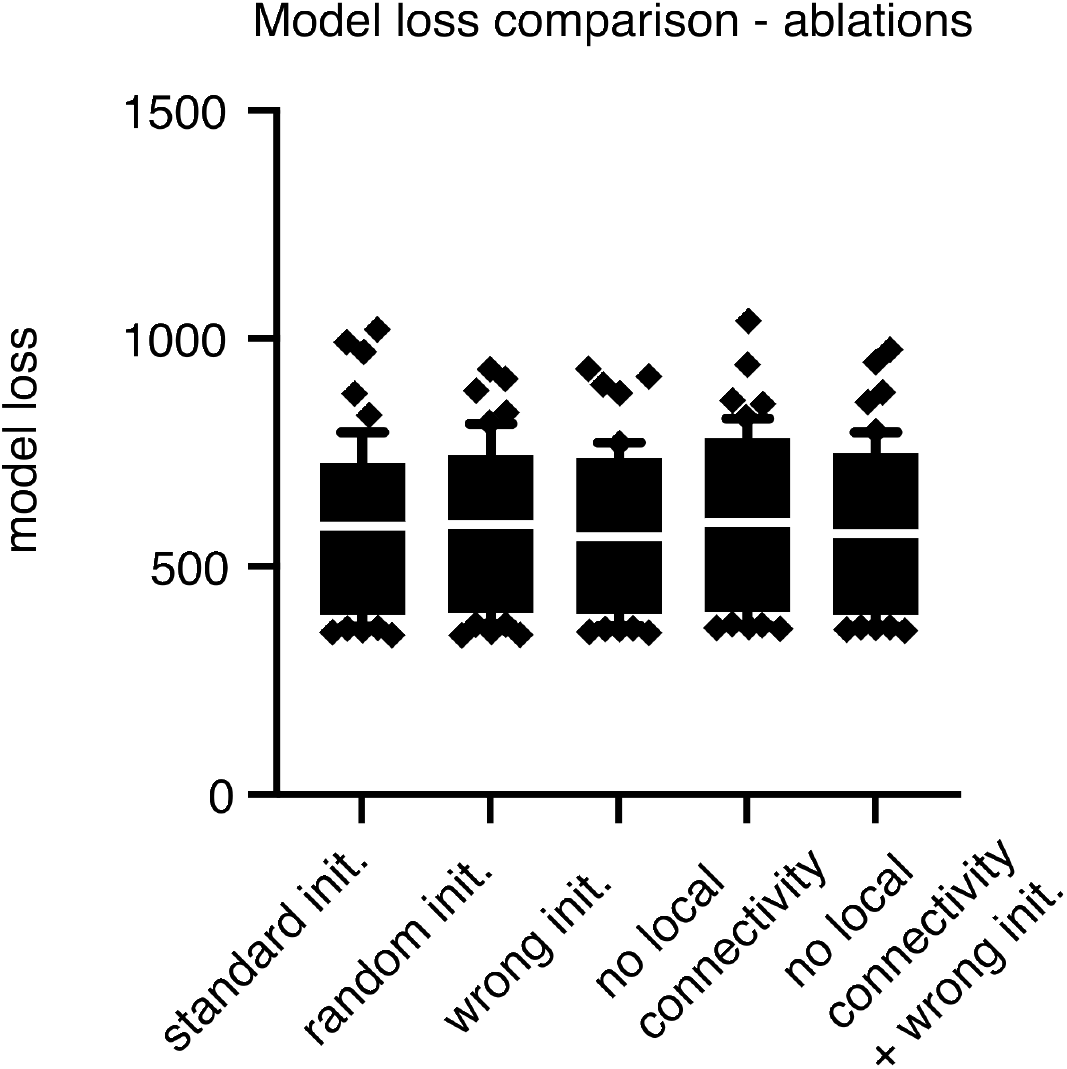
Same plotting conventions as Fig. 5e.

### Supplementary Note 6: Reduced trial-weighted DDMs

**Supplementary Figure 6.**
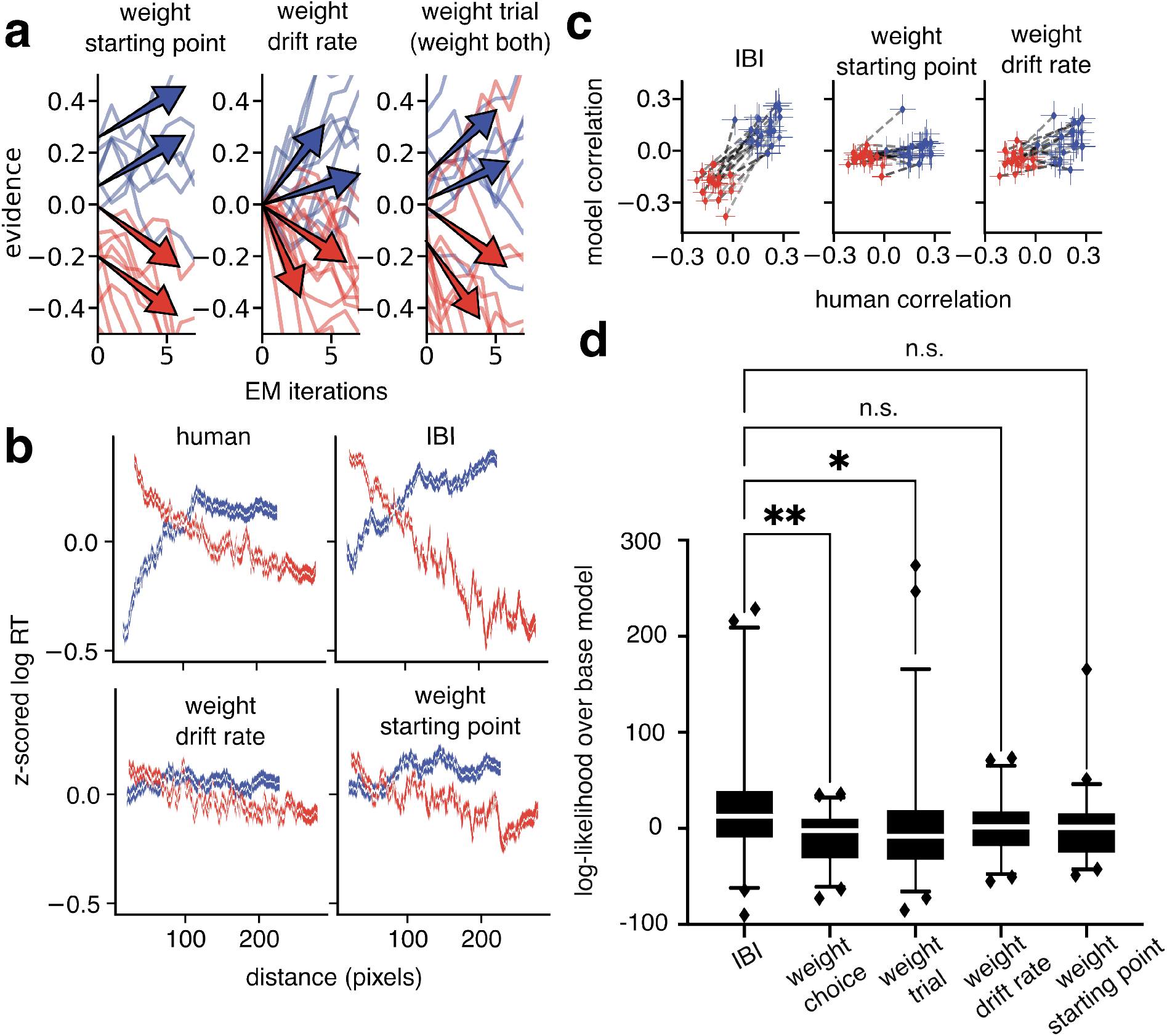
Reduced trial–weighted DDMs do not capture human correlations. **a**, Same plotting conventions as in Fig. 4a. **b**, Same plotting conventions as in Fig. 4c. **c**, Same plotting conventions as in Fig. 4d. **d**, Same plotting conventions as in Fig. 4e.

See Supp. Note 15 for a mathematical description of DDM assumptions. Reduced trial–weighted models are those in which only one of *z* or *m* from Supp. Note 15 eq. S49 is weighted. Weighing only *z* is referred to as the weight starting point model and weighing only *m* is referred to as the weight drift rate model. Correlation plots in Supp. Fig. 6 are flatter than the corresponding plot for the weight per–trial model in Fig. 5d.

### Supplementary Note 7: Per-case model comparison

**Supplementary Figure 7.**
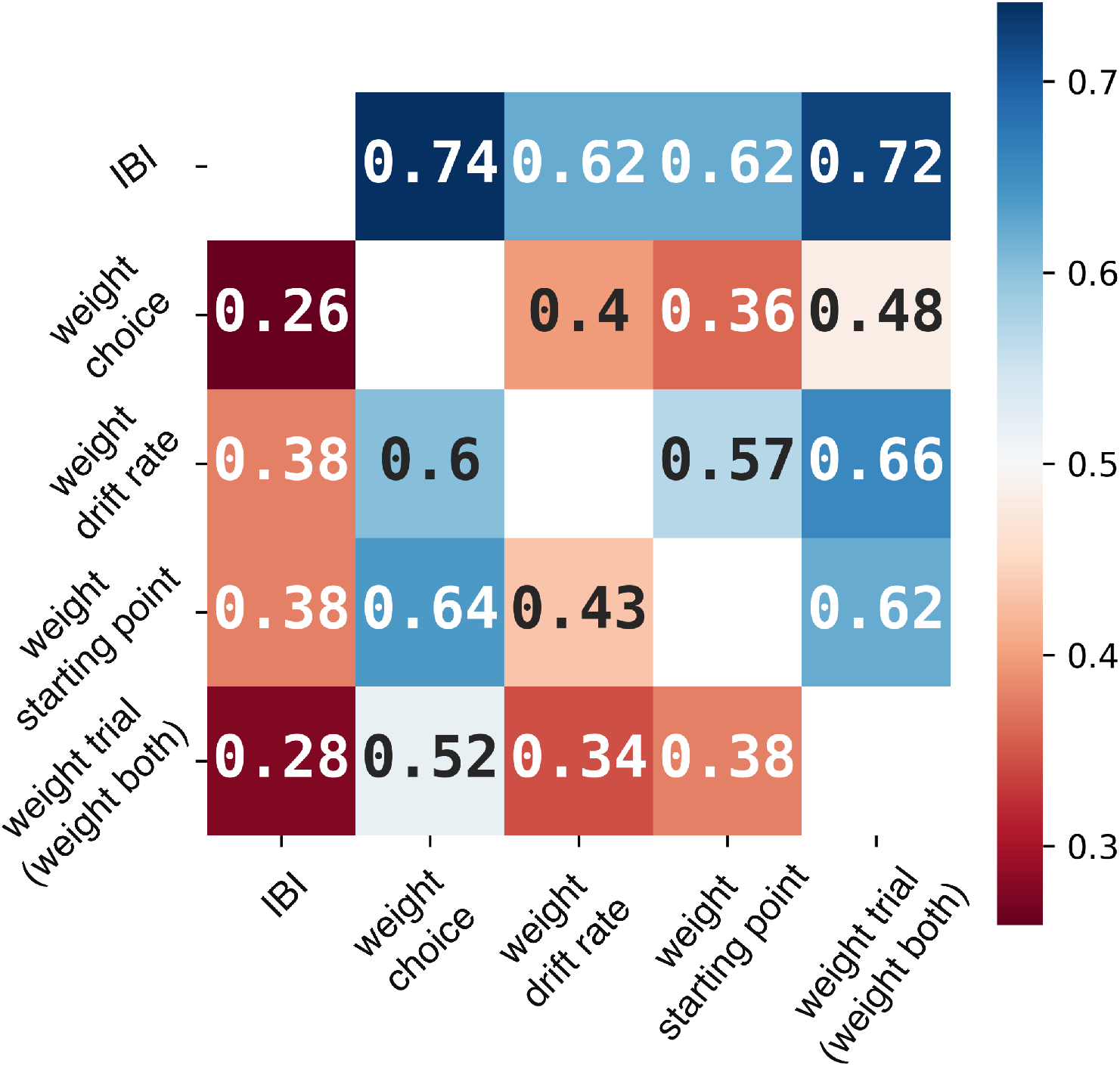
Model comparison, all cases. Annotations in each cell indicate the proportion of cases in which the model specified in the row label is better than the model specified in the column label (e.g. IBI is better than choice-weighted DDM in 74% of cases, out of 58 total.)

### Supplementary Note 8: Bias against *κ*

**Supplementary Figure 8.**
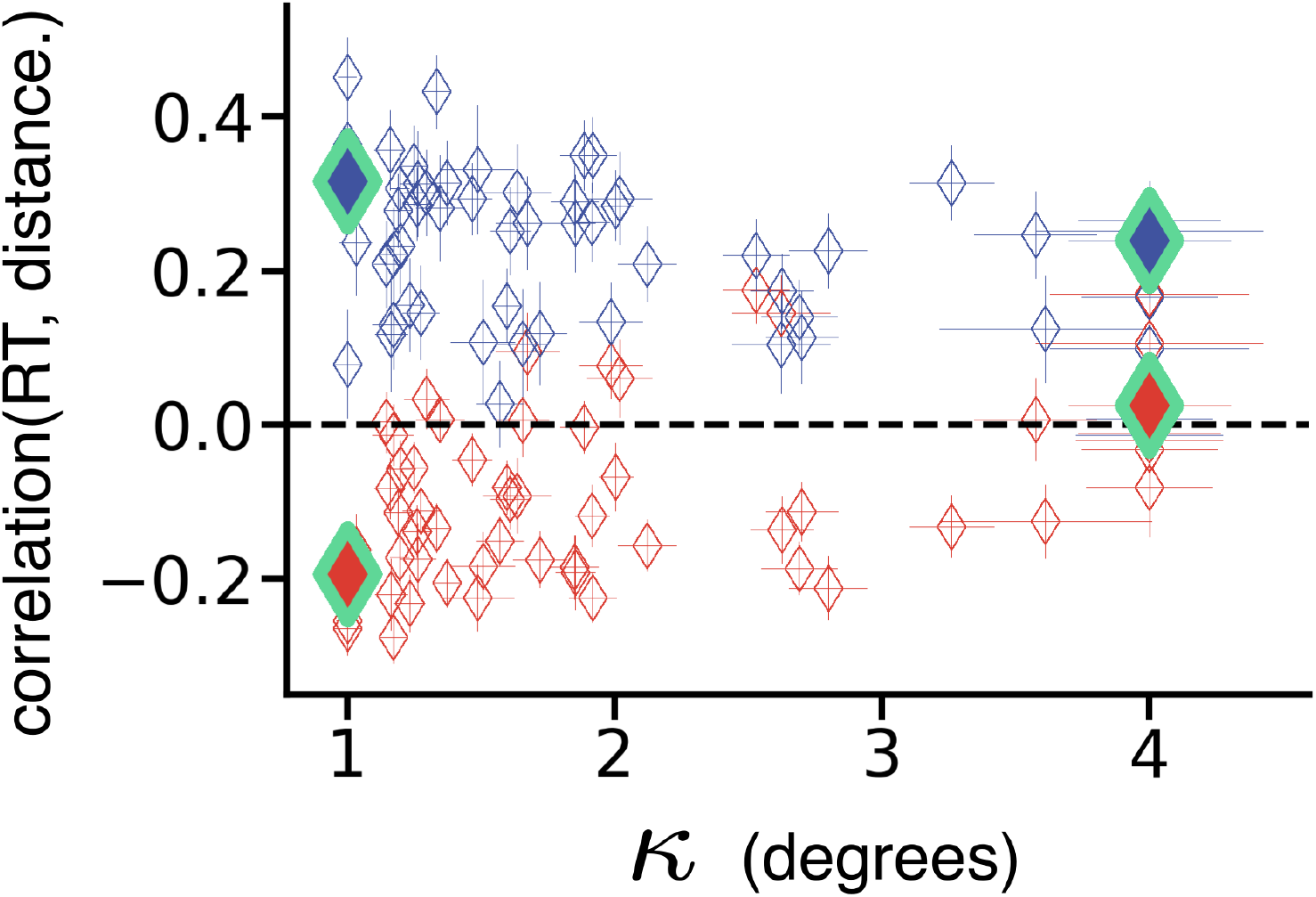
A stronger bias leads to higher RT-distance correlations. Same as Fig. 7d, as a function of the space constant of the exponential fit, *κ*, instead of a function of the amplitude *A*. A smaller value of *κ* corresponds to stronger bias. Green diamonds refer to the same examples as in Fig. 7c.

### Supplementary Note 9: Results without trial exclusion

**Supplementary Figure 9.**
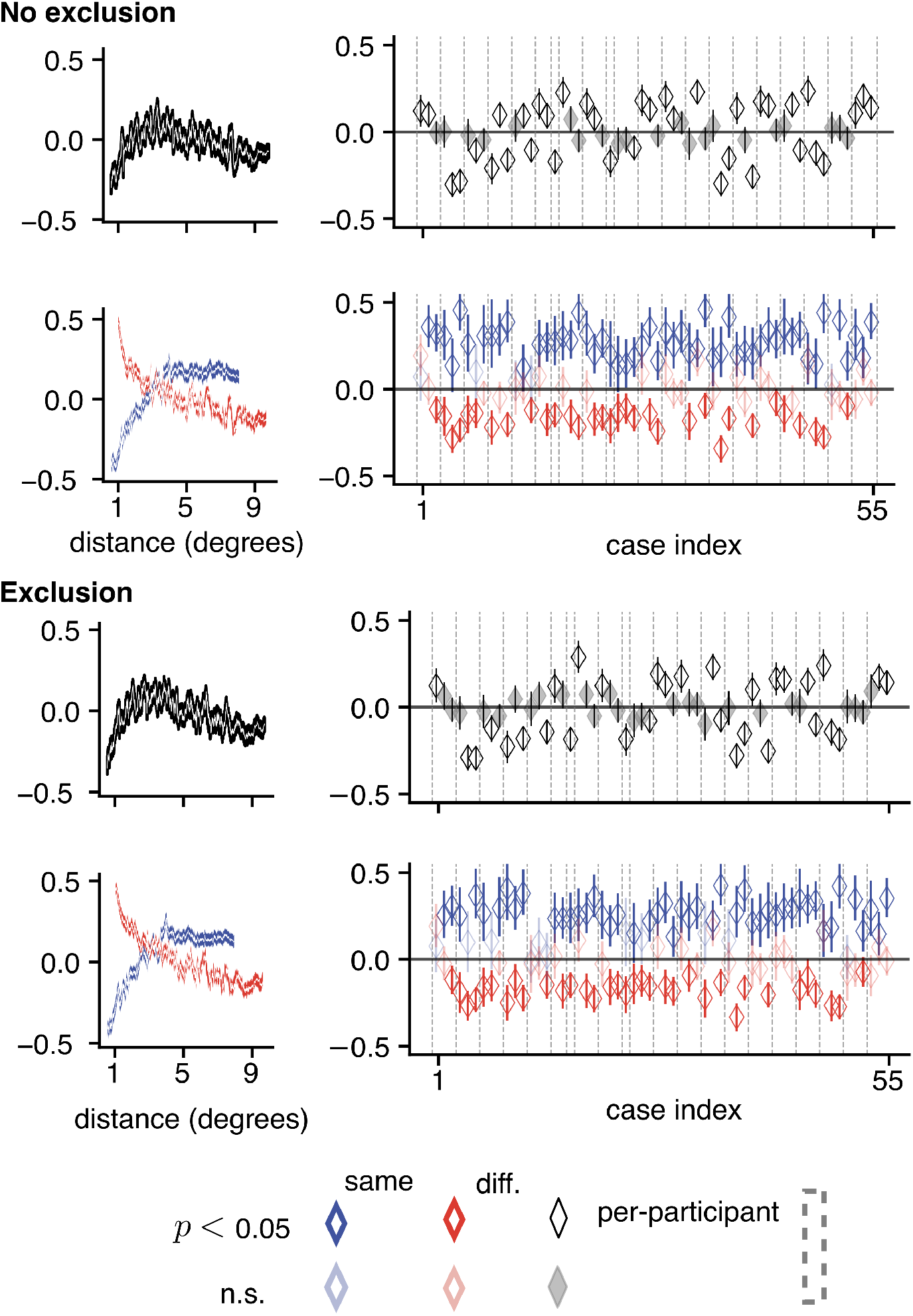
Trial exclusion does not affect results. Top, same as Fig. 2, but without excluding the trials with the 10% slowest RT as detailed in Methods. Bottom, reproduced Fig. 2.The results are qualitatively indistinguishable.

### Supplementary Note 10: Comparison of model performance with different decision rules

**Supplementary Figure 10.**
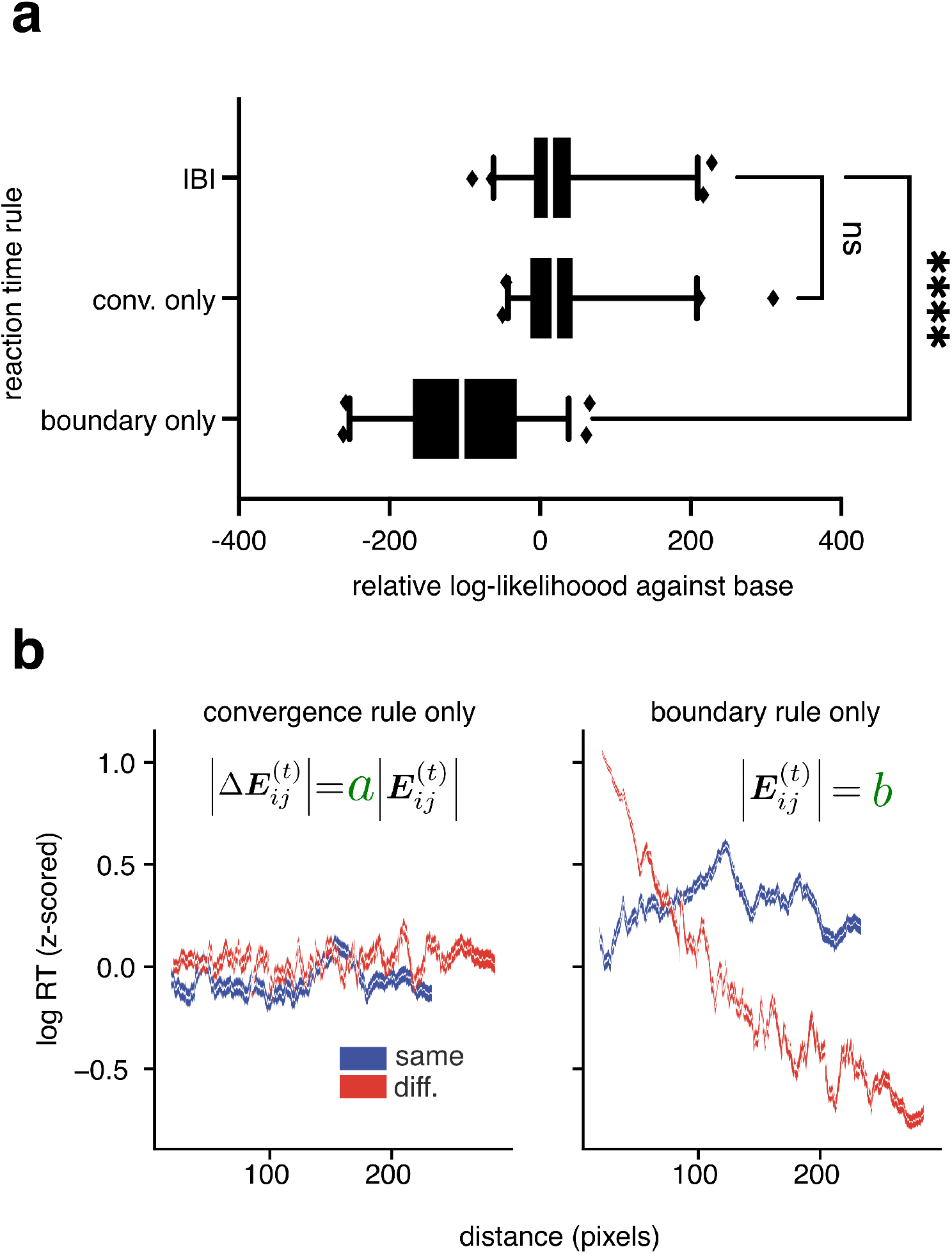
Comparing decision rules. **a**, Same plotting conventions as Fig. 4e. **b** Same plotting conventions as Fig. 3f.

To implement the convergence rule, we smoothed the evidence using a small sliding window (3 iterations) when taking the derivative. We also ensured that convergence was not transient by prescribing that the rule needed to be followed for at least 3 iterations.

The convergence rule is largely responsible for matching model distributions to human RT distributions (Fig. 10a). However, we can observe that the match to human time-distance correlations arises only when a boundary is used for the decision rule (Fig. 10b), implying that natural scene segmentation decisions involve a threshold on certainty.

### Supplementary Note 11: Spatiotemporal trends in consistent vs. inconsistent responses

**Supplementary Figure 11.**
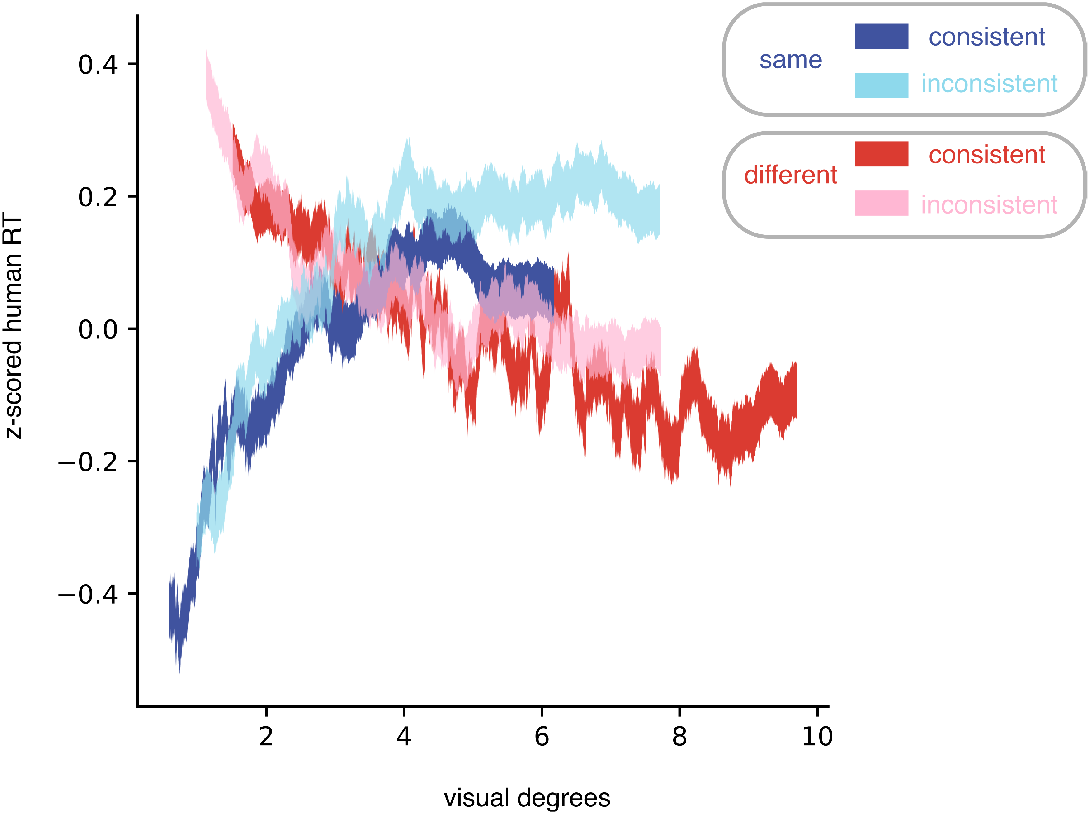
Same plotting conventions as Fig. 2b, without the white mean lines.

### Supplementary Note 12: Cases with stronger spatial bias are better fit with IBI than DDMs

**Supplementary Figure 12.**
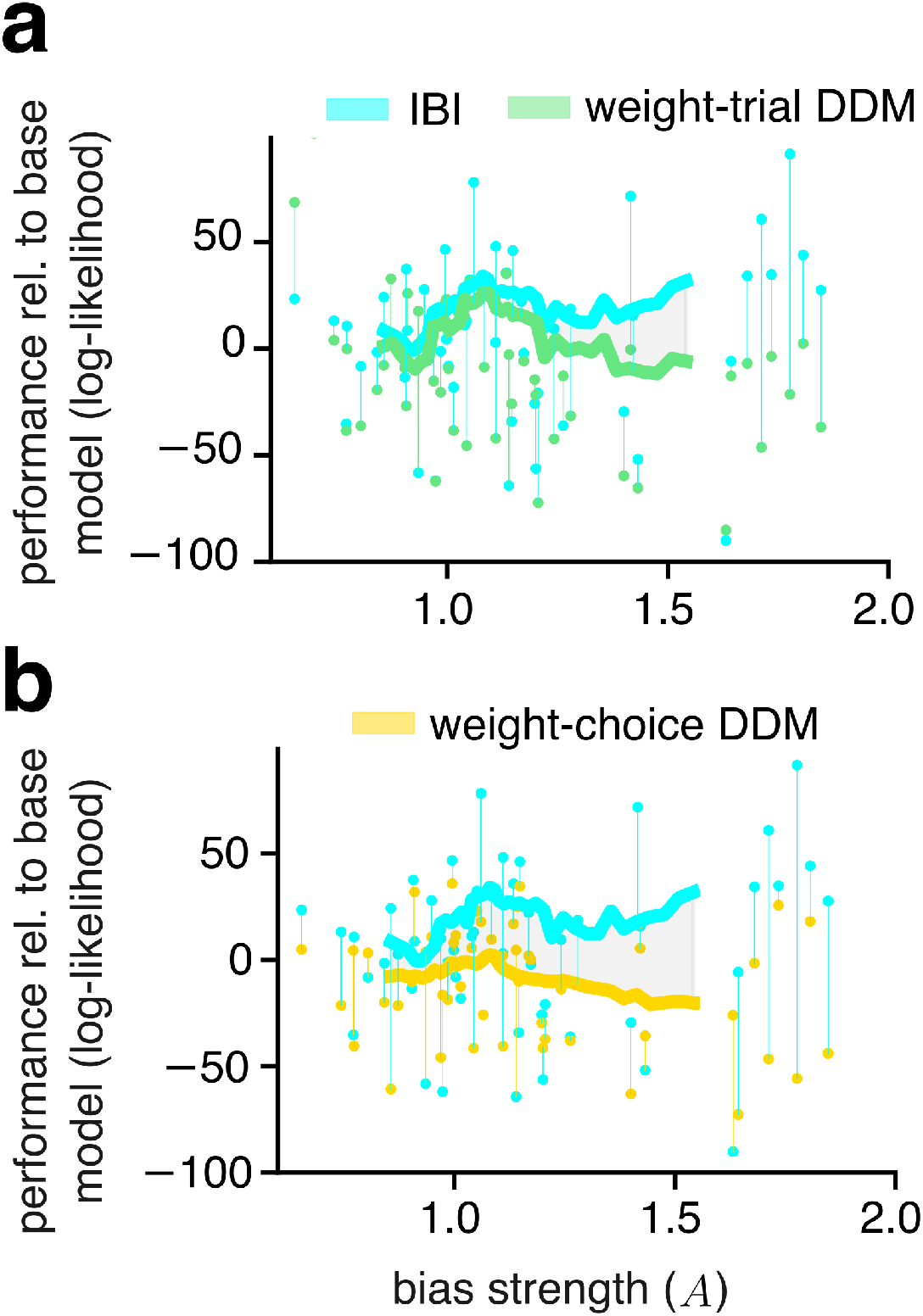
Cases with stronger human bias distinguish models with versus without spatial prior. **a**, Cross-validated model performance relative to the base model (defined as in Fig. 5e) as a function of bias strength. Each dot denotes model performance for one case. Each vertical line connects the performance of the IBI model (cyan) and a trial-weighted DDM (green). Thick lines: running average of the models performance with a sliding window of 15 cases. The shaded gray area indicates the difference in average performance between the two models. **b**, same as panel **a** but comparing IBI versus the choice-weighted DDM. In **b**, 6 cases with data points outside *y*-axis limits were excised to improve readability (but were included in the running average).

### Supplementary Note 13: Reaction times decrease throughout the experimental block

**Supplementary Figure 13.**
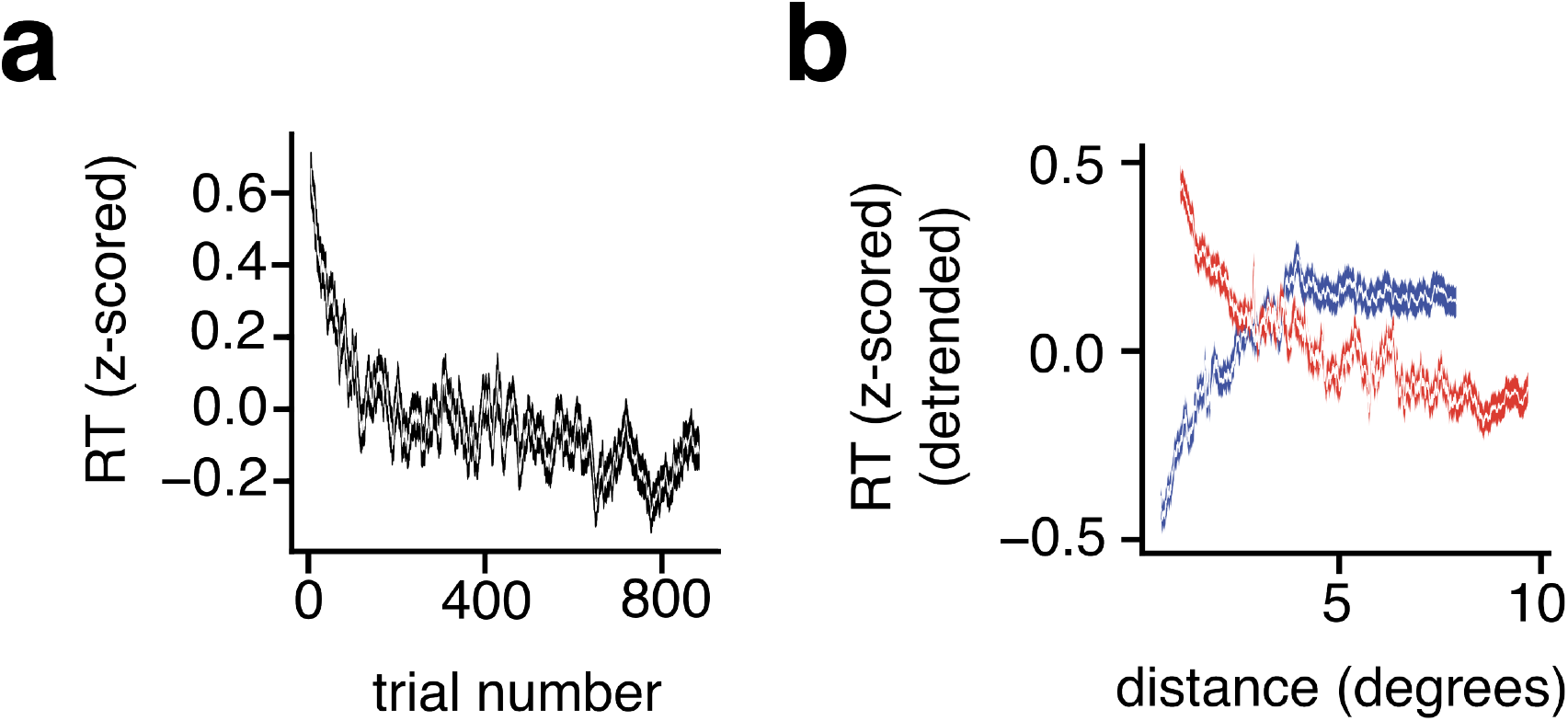
**a**, *z*-scored reaction time vs. ordered trial number. Trials are aggregated across all cases. The white line represents the mean within a sliding window of 600 trials. Black shaded area: s.e.m. **b**, Same as Figure 2a, bottom, with detrended data (*i*.*e*. removing the mean trend of panel **a**; details in the text below).

We observe that RTs decrease as the task goes on (Supp. Fig. 13a). To ensure that the presence of this trend (which is unrelated to the distance between cues) did not affect the RT correlations with distance, we detrended the RT data and verified that results did not change. To detrend the data, we fit a decaying exponential to the z-scored RT vs. trial number, and then subtracted the predictions made by this exponential from the empirical z-scored RT values. Supp. Fig. 13b shows that this detrending procedure did not change our findings.

### Supplementary Note 14: Mathematical details

In this section, we derive the update rules presented in equations 1,2, and give an overview of the normative theory underlying our model (subsections A-D). We also discuss reduced versions of the IBI model (subsection E), and how the model produces evidence for a “same segment” or “different segments” decision (subsections F,G).

#### A. Feature extraction

We begin with the sensory input, an RGB image **I** with height *h* and width *w* (in pixels). The first step is to extract features from the RGB image, which we accomplish using the deep convolutional architecture VGG-19 ^2^. Deep convolutions remap RGB values onto high-dimensional feature vectors that have been shown to be useful as predictors of neural activity in early cortex ^3^. Deep neural networks have also been shown to align with the hierarchical human visual system ^4^. In principle however, any shallow or deep feature extraction applied to RGB images can be used with the probabilistic inference used in the steps described in later subsections.

Notation:

- **I** *∈* ℝ^*h×w×*3^ : **I** is a tensor of shape (*h, w*, 3) *e*.*g*. an RGB image.
- *l ∈* [1, 16] : the layer of the neural network.
- **A**^[*l*]^ : the activation from layer *l* of the neural network.
- **W**^[*l*]^ : a square sliding window kernel with size 3 and feature dimension *F*^[*l*]^, at the current layer. E.g. for layer 1: *F*^[1]^ = 64, **W**^[1]^ *∈* ℝ^3*×*3*×*3*×*64 2^.
- 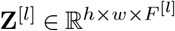 : the output of convolution operation
- 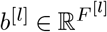 : the bias at a given layer

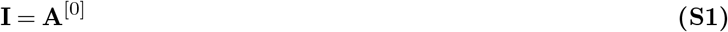

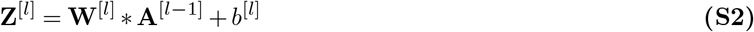

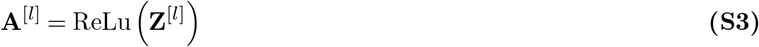

The above equations describe linear-nonlinear transformations of the input as it is passed through sequential layers. For all layers *l, \**is the discrete 2D convolution operator. Values for *W* ^[*l*]^ and *b*^[*l*]^ constitute filters in an *F*-dimensional space, and are learned by training on the ImageNet1K database ^2^ for object recognition (not segmentation). We use the pre-trained weights from ^2^.

For each layer [*l*] we use a linear transformation, principal component analysis (PCA), to reduce the dimensionality of the observations ^5^ to *M*. We set *M* = 6 to standardize feature dimensionality across layers, and because it typically explained 95% of the variance in the shallowest layers. The projection onto principal components is a matrix multiplication:

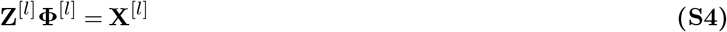

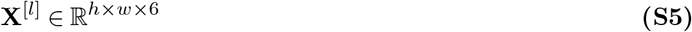

where principal components are the columns of **Φ**^[*l*]^. Therefore, **X**^[*l*]^ is a 6-dimensional representation of the features at layer *l* in image data.

As in the original FlexMM implementation, to improve segmentation performance, we concatenated the low-dimensional representation at any given layer to the PCA projection of RGB features **X**^[0]^. We then defined the feature vector for a pixel, *i*, at layer *l*:

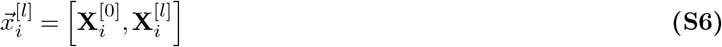

##### A.1. Feature selection

For each image, we selected, out of the first five convolutional layers, the layer whose features led to model segmentation maps most similar to the participant’s segmentation maps decoded from the task (with similarity measured using the adjusted Rand Index ^6^). Therefore the layer of VGG-19 used was tuned to each image for each participant.

We found that intermediate layers of VGG-19 led to results best aligned with human RT-distance correlations. When features were selected from layers deeper than layer 5 or from only layer 1, model correlations were weaker. It is thought that early layers of VGG-19 have access to bottom-up image feature information such as edges, texture, color, and luminance ^2^. Furthermore, because VGG-19 is a network trained for object recognition with backpropagation, deeper layers should have more access to object–specific or semantic information. Therefore, our feature basis partially accounts for semantic or object priors in human participants, but is not fully abstracted away from lower–level image information.

#### B. Gaussian scale mixtures

To simplify our notation we remove the layer–indexing from here on. Bold, italic letters indicate a vector random variable at a single pixel 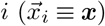 be the random variable representation of a single (*i*.*e*. at one pixel) multidimensional feature vector.

We can define the prior for ***x*** rigorously through normative theories of natural vision. It has been shown that natural images present with the statistics of a Gaussian Scale Mixture (GSM), which is a standard multivariate normal ***g****∈* ℝ^12^ multiplied by a random global scalar *v ∈*ℝ^+^ . The components of the multivariate normal represent the weight with which local features (such as edges with a specific orientation and spatial frequency) contribute to the input image, while the scalar multiplier represents a global modulator such as contrast ^7–11^. In our formulation, we use the Student-t distribution, which is rewritten as a Gaussian Scale Mixture by scaling the covariance of a multidimensional Gaussian, **Σ**, using a scalar *v*, that is itself a random variable sampled from a one–dimensional inverse Chi-square distribution scaled by its degrees of freedom *ν*. Ultimately we write:

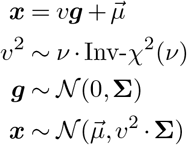

Compute the PDF of ***x***, *p*(***x***) by marginalizing over *v*

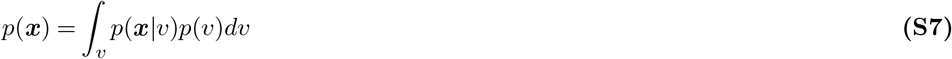

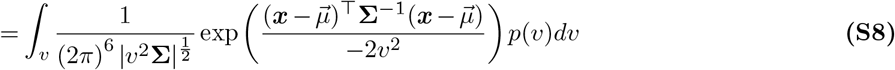

The integral in equation S8 can be written as the expression *f*_1_ ^12^:

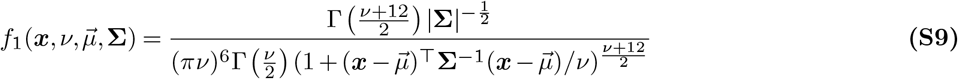

Where Γ is the gamma function. This formulation of the Student distribution makes our parametrization explicit, 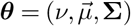.

##### B.1. Mixtures of Gaussian Scale Mixtures

The objective now is to learn a *mixture* of GSMs such that each vector ***x*** may be assigned to one of *K* classes (classes in our formulation are segments). Classes are notated as one-hot encoded vectors and we write *p*(***c***) to refer to the corresponding probability, *e*.*g*. if there were 3 classes, class 2 would be represented as: [0, 1, 0] with corresponding probability *p*(*c*_2_) *≡ p*(***c*** = [0, 1, 0]).

Classes are assigned with a prior probability vector denoted ***π*** which is an element of the *K−*dimensional simplex ***π*** *∈* Δ^*K*^, namely all its elements must sum to one. The PDF of ***x*** is now simply a weighted sum:

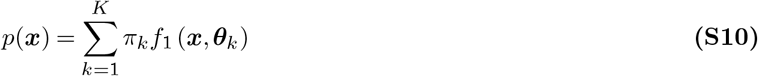

Where *π*_*k*_ is the *k*th-element of the vector ***π***, and ***θ***_*k*_ is the set of parameters 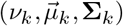 that is unique to the *k*-th class.

#### C. Defining the graphical model for FlexMM

In Supp. Fig. 14, we define our mixture of GSMs as a graphical model, specifically a Bayesian Network, where for two random variables ***r***_1_ and ***r***_2_, the notation ***r***_1→_ ***r***_2_ indicates a conditional relationship *p*(***r***_2_ |***r***_1_) ^13^. Following the FlexMM family of probabilistic mixture models ^5^, to introduce the spatial prior, we add extra edges to the graph (Supp. Fig. 14, blue lines). These edges ultimately define a latent variable ***B****∈* ℝ^*K*^, the concentration parameter of the Dirichlet distribution of ***π***. Only one ***B*** node is shown in Supp. Fig. 14 for the central pixel in a 3 *×*3 neighborhood of pixels. However, there is a ***B*** node for every pixel which is influenced by that pixel’s neighbors.

The extra edges inserted into the model cause a loop. Learning and inference can be performed despite the presence of the loop, as we will discuss in the next section.

**Supplementary Figure 14.**
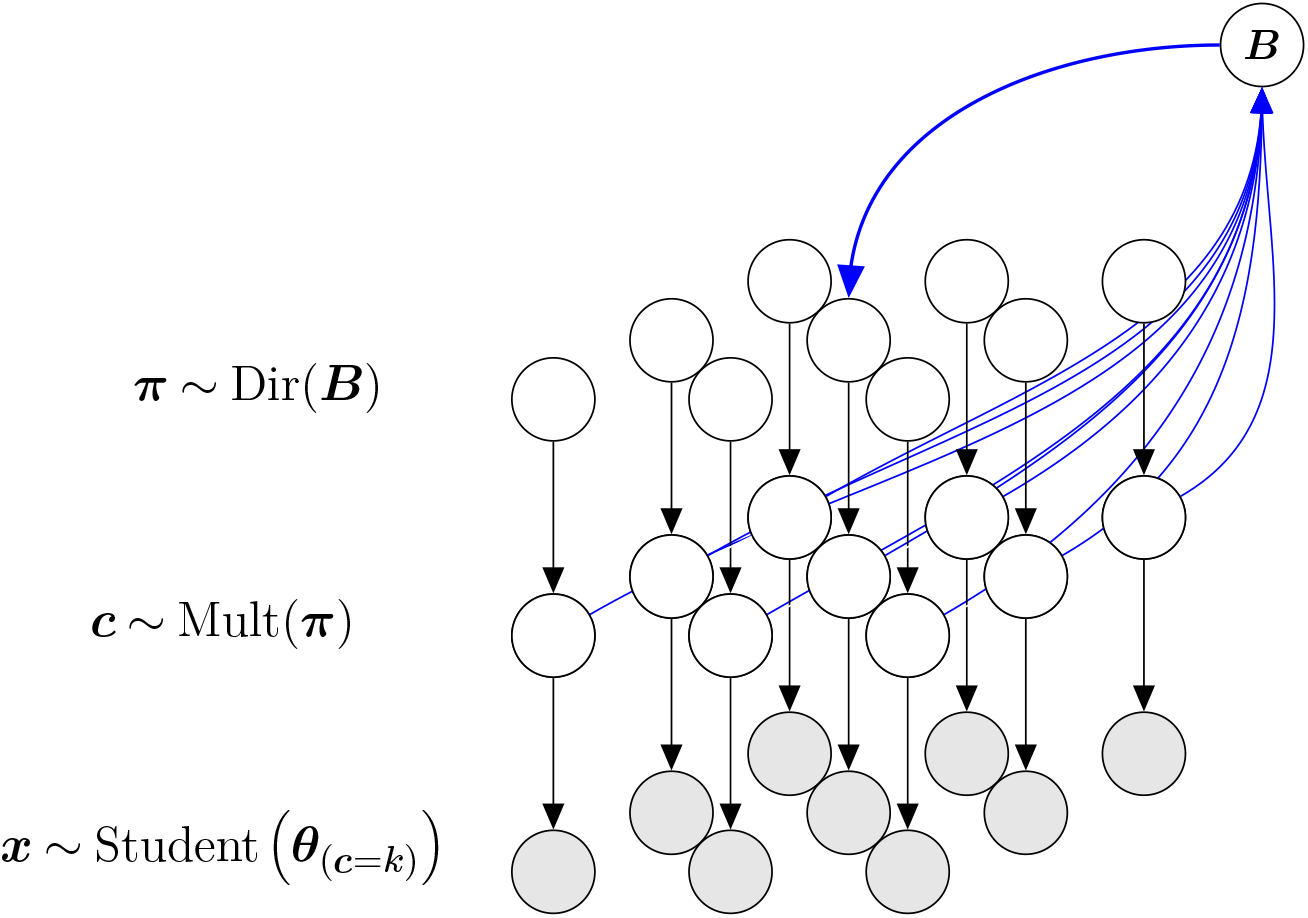
A probabilistic graphical model for a 3 *×* 3 pixel input. We define observed random variables as ***x***, these variables are distributed according to a Student distribution with parameters *{****θ****}*_***c***=*k*_. The parameters are determined based on feature classes ***c***. These prior probabilities of these class labels are defined by the random variable ***π***, distributed according to a Dirichlet distribution. A neighborhood (which is 3 *×* 3 pixels, in this illustration), of surrounding class labels defines the hyperparameter ***B*** (only one ***B*** is shown) which is the concentration parameter of the Dirichlet distribution. Defining ***B*** on the basis of the class labels of surrounding pixels ensures that ***π*** variables that are spatially close to each other are more likely to be similar.

#### D. Learning and inference dynamics

The main goal of this section is to provide a derivation of equations 1,2 of the main text, which our results are based on. As with any probabilistic graphical model, there are multiple algorithms that can be considered for learning and inference ^13^. For FlexMM ^5^ we use a modified version of the Expectation-Maximization algorithm ^14^ that guarantees closed-form linear updates for the class probabilities and flexibility in how class information is integrated across pixels to define the ***B*** variables. The implementation and proof of convergence for FlexMM can be found in our earlier work ^5^.

In the following sections D.1 and D.2, we focus on alternative mathematical views of our model’s dynamics in two way to highlight links to normative theories of perceptual dynamics. First, we briefly mention the recurrent message-passing dynamics that are present throughout the lattice of pixels to connect our work to normative theories of recurrence. Second, we use a view of EM as variational approximate inference to show that iteration is used to minimize statistical free-energy, which leads to a simplified derivation of the update rule in equations 1,2.

We note that disentangling learning (*i*.*e*. estimating parameters given data) from inference (*i*.*e*. forming the posterior distribution, given the data and parameters) requires some care for our graphical model. FlexMM alternates between an M step in which it learns parameters, including a prior probability for the class labels, and an E step in which it infers the posterior class probabilities given those parameters. In the model with the spatial prior (e.g. Fig. 3c and Supp. Fig. 14), the parameters ***π*** are learned per-pixel, so the inferred posterior probability is proportional to the prior. In models without a spatial prior (e.g. Fig. 7f), a single prior probability for all pixels is learned during the M-step, and inferring a posterior probability over segments means that only the E-step generates a probabilistic segmentation map. We will discuss this distinction in greater detail in Section E when discussing the reduced versions of IBI.

##### D.1. EM as recurrent message passing

The Bayesian Network (Supp. Fig. 14) can be rewritten per-pixel with constraints on the relationships between variables. This formalization is called a factor graph ^13^. Supplementary Figure 15 is a per-pixel version of Fig. 3a and Fig. 14 formalized as a factor graph. The factor graph depicts informational relationships with undirected edges, random variable nodes *V* and “factor nodes”, ℱ. Factor nodes ℱ specify the conditional distributions between the variables they connect. The joint probability distribution of the nodes *V* is then written as the product of factors: 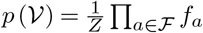, where *Z* is a normalization constant. Specifically, we use the following factorization:

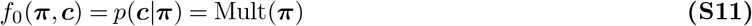

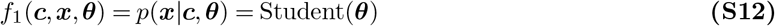

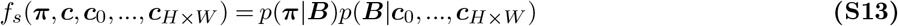

**Supplementary Figure 15.**
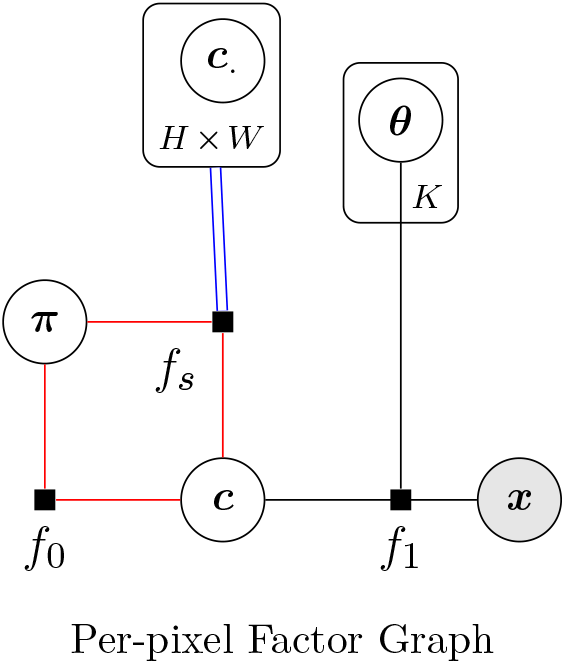
A factor graph of the probabilistic dependencies influencing each pixel. Compared to the probabilistic graphical model of Supp. Fig. 14, directed edges have been replaced with undirected edges and factor nodes. Rounded rectangles around nodes are referred to as “plates” and the annotation within the plate indicates how many values of that random variable are present. Specifically, there are *H × W* neighboring class labels used in the per-pixel class and *K* sets of parameters. The red lines indicate a loop in the graph. To simplify, the random variable ***B*** is not shown as a node, but is implicitly accounted for through factor *f*_*s*_ (eq. S13). The double blue lines indicate that information from multiple neighbors in used in the spatial prior *i*.*e*. ***c*** values over a local set of *H × W* pixels regularize the value of ***π***. *f*_0_, *f*_1_, and *f*_*s*_ are factors of the joint probability distribution 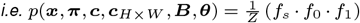.

Equation S13 describes how a spatial prior of size *H× W* (where *H≤ h* and *W ≤w*, the full image size) is implemented where ***B*** is a Dirichlet parameter vector that has the same dimensionality as ***π*** ^15^:

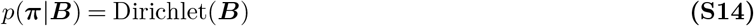

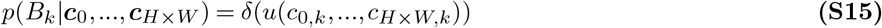

where *u* : ℝ^*H×W ×K*^*→* ℝ^*K*^ can be any linear function, and *δ* is the Dirac delta function.

Our factor graph is written with a loop (Fig. 15), but previous work using the EM algorithm on loopy factor graphs ^16^, has shown that the Expectation and Maximization steps can be considered as message-passing along two separate “tree–like” graphs, and that if neither of them contains loops the EM algorithm can be implemented exactly (see Dauwels ^17^ for the explicit form of messages used in EM). This is the case for our model ^5^ (Supp. Fig. 16).

One advantage of describing FlexMM in terms of message-passing is that it highlights the connection of FlexMM to normative and mechanistic theories that used this powerful computational scheme to clarify the role of recurrence (both lateral recurrence and feedback connections) in probabilistic neural computations ^18–20^.

**Supplementary Figure 16.**
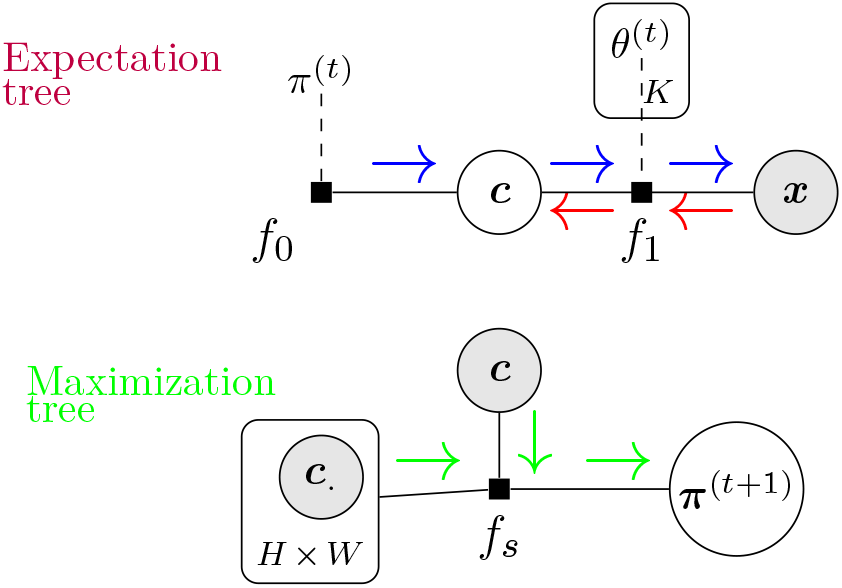
Breaking the loopy model using EM. The Expectation tree (top row) and the Maximization tree (bottom row) generated by “unrolling” the loopy model. Arrows represent messages passed along the edges. *π*^(*t*)^ and *θ*^(*t*)^ are known values of parameters at the current time step and therefore not written as variable nodes here. The algorithm for computing the expectation in the Expectation Step is a forward-backward sum-product message-passing algorithm ^17^. The blue arrows show the forward pass while the red arrows show the backward pass. In the Maximization tree, the spatial prior is implemented using max-product message passing with messages represented by green arrows ^17^.

##### D.2. EM as variational inference

Having discussed the view of EM as a message–passing algorithm that connects to theories of recurrence, we now present an alternative view of EM iterations as approximate variational inference ^15,21–23^. This view emphasizes that iteration in EM is a form of minimizing statistical free-energy, and can in principle, be seen as gradient descent where information from the prior helps determine the direction of optimal descent ^23^. We will use this view to present an abridged derivation.

Implementing the EM algorithm as variational inference involves treating ***π*** and ***θ*** as parameters in order to learn the marginal distribution *p*(***x*** |***π, θ***) that best explains the observations ***x*** at all pixels. Here, we will not explicitly discuss the role of the Dirichlet as a spatial prior, but focus on dynamics. For a set of observations **X** over all *h× w* pixels, we begin with the assumption that a single observation ***x***_*i*_ is independent from others. In this section, the spatial index *i* is explicitly written.

The log-probability of the observations factorizes as follows, where **Π** and **Θ** are the set of mixing weights and Student parameters respectively, for all pixels :

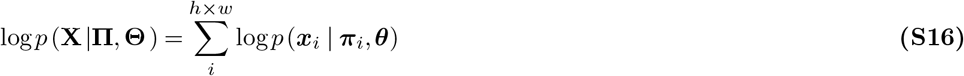

We now introduce the latent variable ***c***_*i*_, a class label, for each observation at pixel *i*. As in earlier sections, classes are notated as one-hot encoded vectors, *e*.*g*. if there were 3 classes, class 2 would be represented as: [0, 1, 0] with corresponding probability *p*(*c*_*i*,2_) *≡ p*(***c***_*i*_ = [0, 1, 0]). Then:

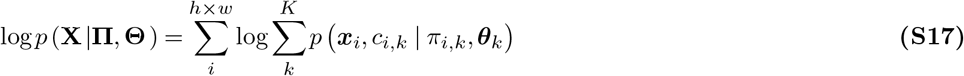

Equation S17 defines the observation likelihood as the marginal likelihood over ***c***_*i*_. To move the summation outside the logarithm we can introduce an approximate distribution for ***c***_*i*_, namely ***c***_*i*_ *∼ q*^(*t*)^.

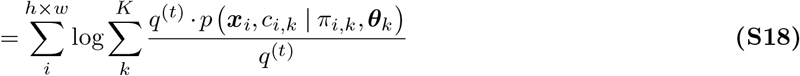

The above can be rewritten using the expectation operator 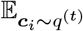, which is the expectation with respect to the approximate distribution *q*^(*t*)^. Using the expectation operator:

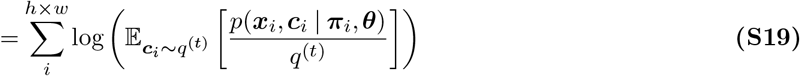

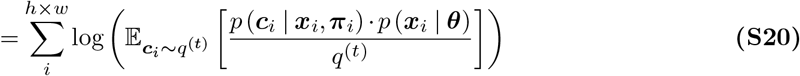

Using Jensen’s inequality we can move the logarithm inside the summation, and establish a lower bound to maximize.

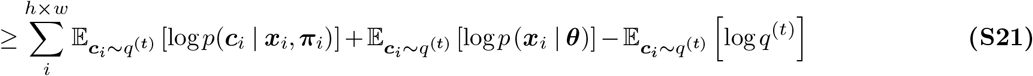

Inference aims to find the approximate distribution *q* to maximize the above lower-bound monotonically. This is equivalent to minimizing the negative of this quantity, which can be interpreted as a quantity called variational free energy ^21,22,24^ (note that the third term is the statistical entropy of the approximate distribution *q*^(*t*)^).

Note that the challenge with performing inference with the model as written in equation S21 is that in the first term:

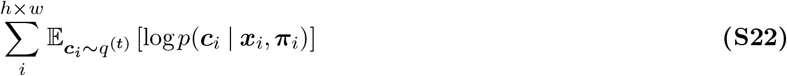

there are (*K −*1) *·h· w* parameters and only *h ·w* observations. Therefore, to effectively reduce the number of parameters we use regularization: we introduce a linear function *u* that can be applied over the ***π***_*i*_ within a neighborhood of size *H× W* to facilitate spatial smoothing. In practice, special care must be taken to define the *u* as linear using additional parameters from conjugate priors (see previous Section D.1 and the FlexMM paper ^5^, particularly Appendix B for proofs). We ignore these additional parameters in the current derivation to focus on one main point — the *u* function must be linear as applied to the expectation so that it can be taken out of the expectation term by the linearity of expectation:

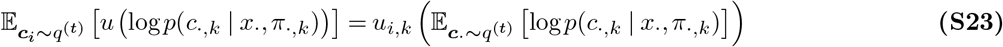

To apply the condition that closer pixels are more likely to be in the same class we use as the *u* function a convolution with a discrete 2D Gaussian with finite kernel size.

##### D.3. Per-pixel EM

Notice that the linear *u* function can be applied to the standard formulation of the expected complete-log-likelihood over neighbors. The standard formulation of EM states that parameters evolve to increase the likelihood of the dataset as a batch ^14^. In our case, parameters are per-pixel. Examining the likelihood traces in our model we found that certain pairs of pixels reach their final likelihood value before other pairs. Therefore it is worth a brief look at existing theory on EM with single observations, termed incremental EM, which was shown to be feasible through the variational view of EM ^21^. Neal and Hinton have shown that maximizing log *p*(**X** |**Π, *θ***) is equivalent to maximizing the complete log-likelihood per-pixel when latent observations {***c***}_*∀i*_ are independent. In our formulation strictly speaking, {***c***}_*∀i*_ are not independent because of the *u* function. However, because ***π***_*i*_ are also learned per pixel we can treat {***c***}_*∀i*_ as conditionally independent when ***π***_*i*_ is known for all *i*. This justifies that single pixel updates are unique while increasing likelihood for the whole image.

##### D.4. EM as gradient ascent

Having established that we can apply *u* to the standard complete-log-likelihood, and that this quantity can be optimized per pixel, we can introduce a per-pixel objective functional *O*_*i*_ by defining a parameter space {***π***_*i*_,*}* ***θ****∈ ϑ* and a distributional space *q ∈Q* . The way that iteration enables gradient ascent for this functional is summarized in Table 3, and aligns with normative work exploring dynamic sampling circuits^25–27^. The equation for the per-pixel objective functional at time step (*t*) (ignoring the entropy contribution from equation S21) is:

**Table 2.** Full IBI model vs. ablations, model loss. We summarize values with a bootstrapped 95 % confidence interval around the median across a 5-fold cross validation per image per participant. The output of our custom loss function for unseen data is averaged across all folds and cases. Lower values indicate a better fit. *p >* 0.5 for all comparisons against the main model.

|  | Initialization: Subjective segmentation map | Wrong image map | Random |
| --- | --- | --- | --- |
| Smoothing |  |  |  |
| Markov smoothing | 531 – 639 | 542 – 600 | 556 – 626 |
| No Markov smoothing | 559 – 642 | 553 – 610 | N/A |

**Table 3.**
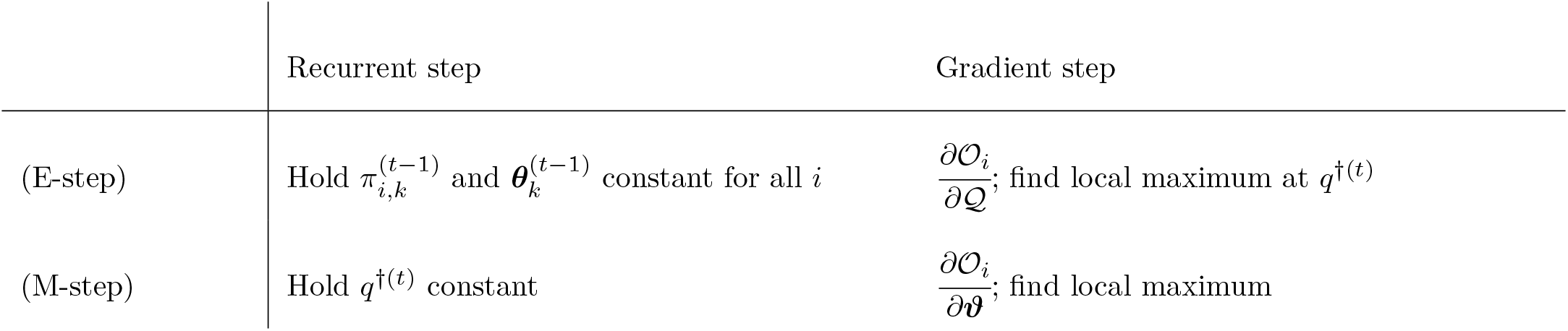
EM framed as gradient ascent ^21,23,28^. This framing shows how different pixels each contribute to increasing the overall data likelihood as long as there is information in *q*^*†*^ about neighbors. The update rule in equation 2 is derived by finding closed-form solutions for local maxima when setting the gradient steps equal to 0 and then applying the function *u*. Values for 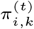 are updated until the total likelihood log *p* (**X** |**Π, *θ*** ) reaches an asymptote.

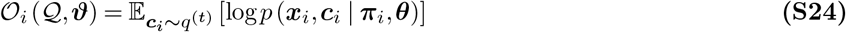

Consider the derivative 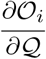. Past work has shown ^21^ that when setting 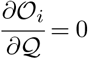 a maximum exists at: *q*^*†*^ = *p*(***c***|***x***). t any given iteration *t*, computing *q*^*†*(*t*)^ = *p*(***c***|***x***) is the expectation step, while the maximization step involves learning parameters 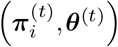 to further improve approximation. The EM update rule as derived below uses parametrized distributions to make these computations exactly, but it is nevertheless a specialized form of gradient-based optimization with the goal of approximation. These closed–form expressions meaningfully capture dynamics because constrained optimization (*i*.*e*. ***π*** must sum up to one) require a small step size. This was shown by Xu and Jordan ^28^ who explicitly relate EM to iterative Newton and Quasi-Newton’s methods.

To continue the derivation, for the M step we re-express the expectation in equation S24 explicitly as a sum over class labels and we introduce explicitly the time labels:

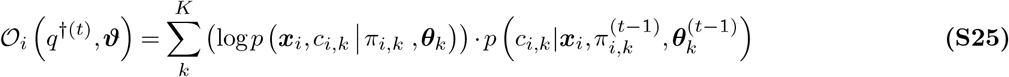

We use Bayes’ rule for the conditional probability term:

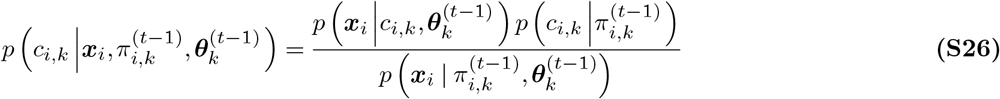

We write the denominator as the marginalized sum over all classes. We also introduce the symbol 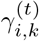, termed the “responsibility” which quantifies how “responsible” a particular class is for a particular observation.

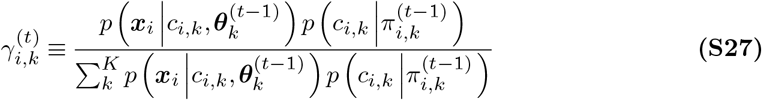

Now we can substitute equation S27 into equation S25:

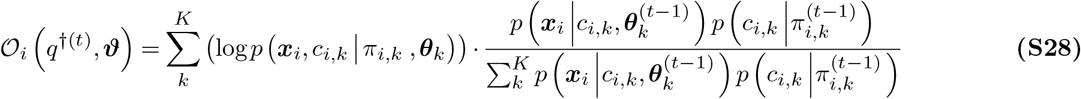

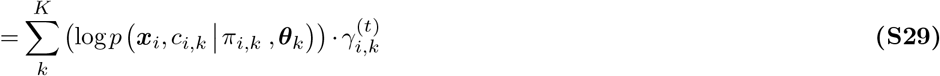

Decompose the joint probability distribution

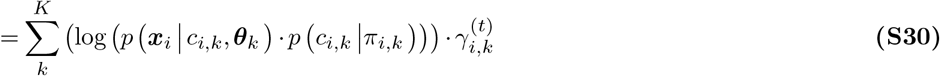

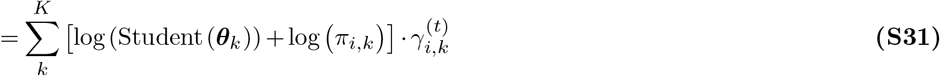

Here we show the maximization with respect to *π*_*i,k*_ (see Peel ^12^ for estimation of the parameters ***θ***_*k*_ ). Maximization for *π*_*i,k*_ is subject to the constraint 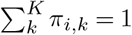, implemented with a Lagrange multiplier *λ*:

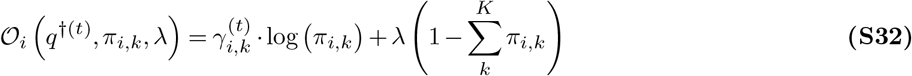

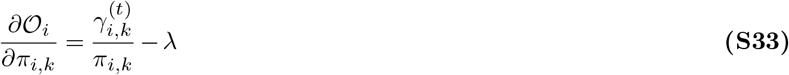

Set the partial derivative equal to 0 and solve for *π*_*i,k*_ to find the updated value 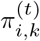:

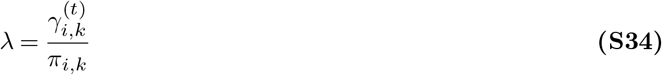

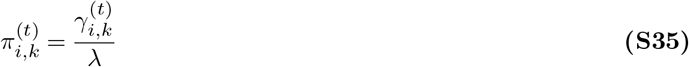

equation S35 is the standard formulation of the EM update rule. Because 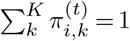 we can write that 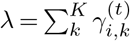. Our formulation differs however because of the *u* function applied to the complete log-likelihood so we write:

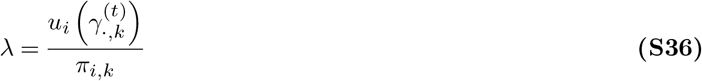

Finally, this leads to our update rule:

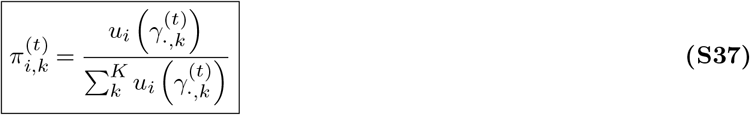

We end with a summary of how our approximate distribution *q* changes with time. We started with an approximate distribution over ***c, c*** *∼q*. We show that to maximize likelihood/minimize free-energy *q→ q*^*†*^ *≡ p*(***c***| ***x***) *≡γ*^(*t*)^ which is then rescaled to *π*^(*t*)^ and that through spatial smoothing with *u*, ***π*** has further, single-pixel dynamics.

##### D.5. Deep spatial prior

As explained in Section A, we use the notation ***x***^[*l*]^ to indicate that our random variable formulation of the observations **X** is in a subspace of **Z**^[*l*]^ for a specific layer [*l*]. This leads to latent variables being layer specific as well (e.g. ***c***^[*l*]^, which was omitted in earlier sections for clarity). Just as class labels ***c*** that are close together in space are more likely to be the same, we assumed that so too should class labels for the same location across neighboring layers {***c***^[*l−*1]^, ***c***^[*l*]^, ***c***^[*l*+1]^ }. This leads to better segmentation performance with FlexMM ^5^.

Therefore, we can write equation S22 for a spatial neighborhood while considering neighboring layers as well:

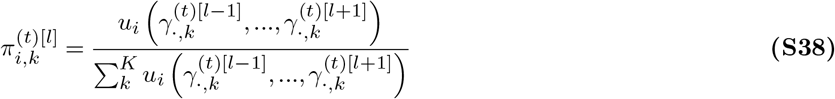

The appropriate linear function *u* for applying this deep spatial prior and the corresponding update rule is presented in Box I of our FlexMM paper ^5^.

##### D.6. EM initial guess

In all versions of IBI and Expectation-Maximization in general, the prior is iteratively updated to generate a posterior and vice–versa. Therefore, technically speaking, the first prior our model learns is based on the initial guess. In the main analysis, the initial guess comes either from participants segmentation maps or from algorithmic maps from DINOv3 features. This initial guess introduces information about perceptually meaningful segments, which are likely based on both bottom-up features as well as object and scene understanding.

For our initial guesses, 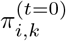 and 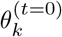, first we define 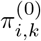 by sampling from resized, probabilistic segmentation maps of the participants (Fig. 1b, top). Resizing is necessary because the coordinate system for participant segmentation maps is lower resolution than the per-pixel coordinate system of FlexMM. Let the value at each pixel in this resized participant segmentation map be notated ***p*** (analogous to ***π*** from FlexMM, that we have discussed in detail, but based on participant responses). We can then define sampled labels:

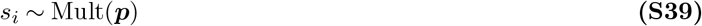

We then compute 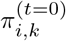 from *s*_*i*_ like so.

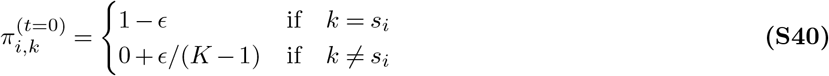

After computing this mapping, we estimate 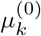 and **Σ**_*k*_ using assignments *s*_*i*_. For the initial guess of *ν* (the scale parameter, see Section B above) we use *ν* = 2, which simplifies the mixer distribution P(*v*) to a scaled inverse Rayleigh distribution ^5,11^.

Note that by sampling in the initial guess, our model is sensitive to different levels of perceptual uncertainty in each case (*i*.*e*. each participant and image). This is part of the reason we refer to the EM initial guess as the participant’s mental map in the main text. While the final maps from FlexMM are different depending on the precise sample of initial guess, our main findings on dynamics are qualitatively robust across varied initial guesses.

#### E. Graphical model perturbations

In this section we focus on the ablations detailed in Figure 6.

##### E.1. No local connectivity model

If we define the *u* function as a Gaussian with an infinitely wide kernel 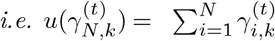, where *N* = *h × w*, then we can plug back into equation S27 to get:

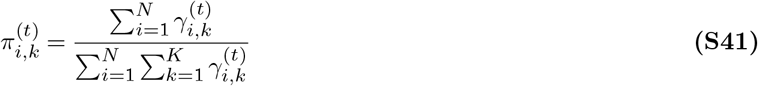

By definition, 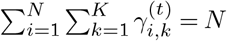 so:

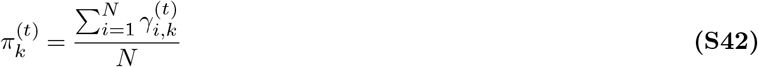

Because we are summing over all *i*, the index *i* is no longer present on the left-hand-side of equation S42. Therefore, to obtain per-pixel probabilities, we have to compute the posterior (which is also equivalent to the E step of the next iteration):

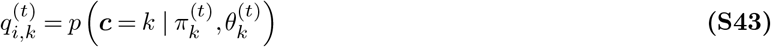

##### E.2. Wrong mental map and random mental map models

For the wrong image initialization we sampled from the participant-image segmentation map that had the lowest match with the standard initialization’s segmentation map (computed with aRI ^6^).

For the random mental map we sampled the initial guess from a multinomial distribution where samples had equal probability of segment label up to the prescribed number of segments *K*:

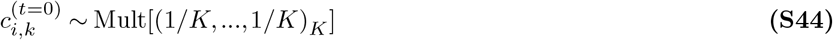

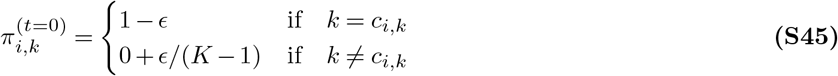

#### F. Model convergence

Having defined the initial state of our algorithm (the “mental map”), and the processing at each time step we now define model convergence.

##### F.1. The iterative timescale

As parameters are updated at each time step, ***π***^(*t*)^ and ***θ***^(*t*)^ can be used to compute the value of the observation log-likelihood per time–step log *p* (**X** |**Π**^(*t*)^, ***θ***^(*t*)^) shortened *L*^(*t*)^. We define the time of convergence, *t*^*\**^

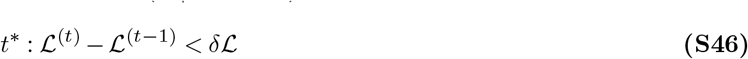

Where *δℒ* is a small, positive constant.

Therefore, we notate the per-pixel segment probabilities at convergence as 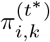. Conceptually, we can think of *t*^*\**^ as the time at which further iteration does not meaningfully increase observation likelihood. In other words, after time *t*^*\**^ there is no informational benefit to the computational cost of iteration.

#### G. Evidence for “same segment”

Thus far, we have described FlexMM which provides us with 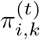. the segment probabilities for each pixel, up until convergence at 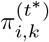. To get from these to evidence about the “same segment” hypothesis, we rewrite equations 3–4 from the main text, Methods.

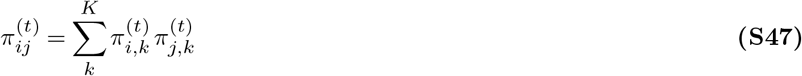

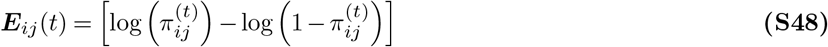

*π*_*ij*_ in equation S47 is the probability that two pixels *i* and *j* belong to the same segment. From this probability we use the log-likelihood ratio as evidence, notated ***E***_*ij*_. Note that although ***E*** is not a vector quantity, we use boldface to emphasize that ***E*** is generated by the IBI model using multi-dimensional inference procedures. The use of the log-likelihood ratio is a principled choice in models of decision–making, including the DDM, as will be further discussed in the upcoming Section 15.

### Supplementary Note 15: Drift Diffusion modeling in this work

#### Drift Diffusion Models

##### Background

We compared the within-trial dynamics of IBI against an alternative with strong support in classical decision-making literature, drift diffusion models (DDMs) ^29,30^. We briefly review the theory before describing the bespoke, image-computable DDMs introduced in this work. (See below for a historical background on DDMs.)

The DDM models an optimal observer—that is an observer who responds the fastest given an acceptable maximum error rate. Sequential samples of evidence, which are corrupted by independent samples of noise, are integrated over time and decisions are reached using the sequential probability ratio test, *i*.*e*. each new sample updates the ratio, *L*(*t*), between the log–likelihood of each alternative hypothesis. DDMs assume that each sample of evidence in a given experimental condition contributes the same amount on average. Based on this assumption, the log–likelihood ratio can be parametrized in continuous time as:

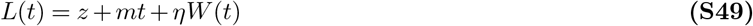

Where *z* is a starting point, *m* the mean rate of evidence (drift) and *W* (*t*) is a Wiener diffusion process such that: *dW∼ N* (0, 1), where the variance of the diffusion process is scaled by *η*. In a DDM, dynamics are unique across trials because drift is perturbed by independent random draws from the Wiener process. The objective of a DDM is not to make per-trial predictions of human choices or reaction times, but to match the characteristically skewed reaction time distributions that arise from perceptual decision-making tasks ^30^.

##### Base DDM

We first fit a base DDM model which was not image-computable. Defining the base DDM through equation S49, we set *z* = 0 and fit *m* directly to the human data per-case without any consideration of image features. *η* was set heuristically and fixed across cases. This procedure is in line with the typical application of DDMs in which the drift rate *m* is not directly encoded in stimulus information, but decoded from fitting to empirical data ^29^.

##### Image-computable DDMs

Image-computable DDMs, hereafter referred to as weighted DDMs, used the segment probabilities from IBI to define the per-trial DDM parameters. In our trial-weighted DDMs we substituted *m* and *z* with the value 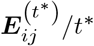 computed from eq. 4, obtained at time *t*^*\**^ which is the time of IBI’s global convergence to a final map. Reduced variants in which we substituted only one of *m* or *z* performed poorly (Supplementary Note 6). In our choice-weighted DDMs, we set *z* = 0 and simply averaged the per-trial values 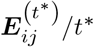 over each choice type grouped by the final map at time *t*^*\**^ (Fig. 5b, middle).

One interpretation of using the IBI evidence at convergence to define the DDM parameters, is assuming that the time required for sensory inference is much less than the time required for integrative decision–making subject to noise (see Supp. 15A.2). In other words, weighted DDMs and IBI capture the same feature and spatial information to use as evidence, but DDMs assume linear within–trial dynamics while ignoring nonlinear dynamics.

#### A. Extended DDM Background

Previous work in decision modeling states that an optimal observer—that is one who responds the fastest given an acceptable maximum error rate—can be modeled using the drift diffusion model (DDM) ^30^. The goal of a DDM is to predict reaction time distributions per choice–type ^31^. In this work, we do fit DDMs to reaction time distributions, but our primary goal is not prediction. Instead we wish to compare the widely-adopted stochastic linear dynamics of a DDM to the nonlinear dynamics of IBI. For fair comparison, we developed an image-computable DDM and here we review the assumptions we followed in order to do so.

##### A.1. Sequential log-probability ratio test

First, we define the fundamental assumptions of DDMs. DDMs model decision-making as a sequential analysis problem, in which evidence, *ξ*, about the stimulus is accumulated over *s* sequential samples (*i*.*e. ξ*_0_, *ξ*_1_, …, *ξ*_*s*_). The optimal way to consider this evidence is using the sequential probability ratio test ^30,32–34^, where *h*_y_ is the hypothesis that two cues are in the same segment while *h*_n_ is the hypothesis that two cues are in a different segment:

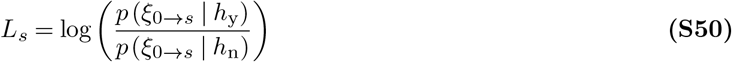

Assuming that samples are independent we can take the limit approaching a differential amount of time (*dt*):

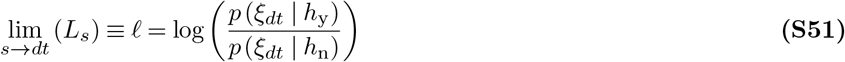

In DDMs, a simplifying assumption is made that because evidence is drawn from a large population of neurons perturbed by noise, the central limit theorem applies, and the differential log–likelihood can be re-parametrized as a Gaussian with mean *m* and variance *η*^2^ (see Bogacz ^30^ Appendix A for a more detailed derivation).

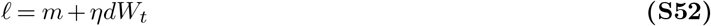

Classical DDMs use a stochastic process called the Wiener process to model noise. The Wiener process *W* (*t*) is introduced such that 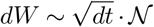 where *N* is a Gaussian with zero mean and unit variance. Following from the independent samples assumption made earlier: for 0 *< s < t < u < v, W* (*v*) *−W* (*u*) is independent from *W* (*t*) *−W* (*s*)^30^. Using the Wiener process we can reparametrize log-likelihood ratio *L* as a function of time *L*(*t*):

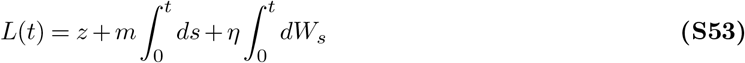

Where *z* is introduced as a starting point bias. Integral notation makes clear that evidence accumulation (a.k.a. evidence integration) refers to averaging out noise over time, but here on we write *L*(*t*) without integrals for simplicity, like so:

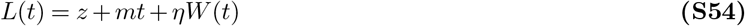

##### A.2. Iterative and Integrative timescales

Treatment of the differential log–likelihood *l* is where FlexMM differs from classical DDMs. In FlexMM, the dynamics of *l* is determined by the information that FlexMM extracts from the image, and thus *l* is affected by the connectivity of the graph. Comparing *L*(*t*) to ***E***_*ij*_(*t*) from equation S48 in Supplementary Note 14, we can appreciate that they are both log-probability ratios. Thanks to this shared probabilistic formulation, we were able to further develop bespoke DDMs that leverage image statistics while eliminating the nonlinear within-trial dynamics of iteration in IBI.

The intuitive approach was to assume that we can substitute the parameters in *L*(*t*) with value of the *unchanging* log-likelihood at iteration convergence ***E***_*ij*_(*t*^*\**^), divided by *t*^*\**^. We introduce a new variable for differential evidence in the integrative timescale:

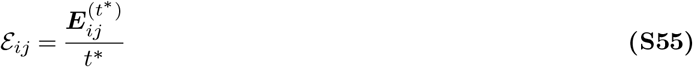

This quotient is the average value of sensory evidence during the iterative timescale. Notionally, we divide by *t*^*\**^ to keep our units consistent with *m*. Practically, this division ensures that stimuli that take longer to converge do not have their evidence count for more, which would be counter–intuitive.

Therefore, technically speaking, *L*(*t*) is only defined for *t > t*^*\**^. Following the assumptions of sequential analysis, this creates a decision-making model in which the integrative timescale follows the iterative timescale. The model which yields a better fit is deemed to “set the tempo”. In other words, it is a better model of the slow step of the decision-making process. That is, if DDM is a better model, extracting the evidence from the input image either takes the same amount of time for all pairs or is much faster (and thus has less influence on reaction times) than the process of averaging out noise. Conversely, if IBI is a better model, reaction times are largely determined by the dynamic process that extracts evidence from the sensory input, whereas additional noise is negligible. See Table 4 for a side–by–side comparison of timescales.

**Table 4.**
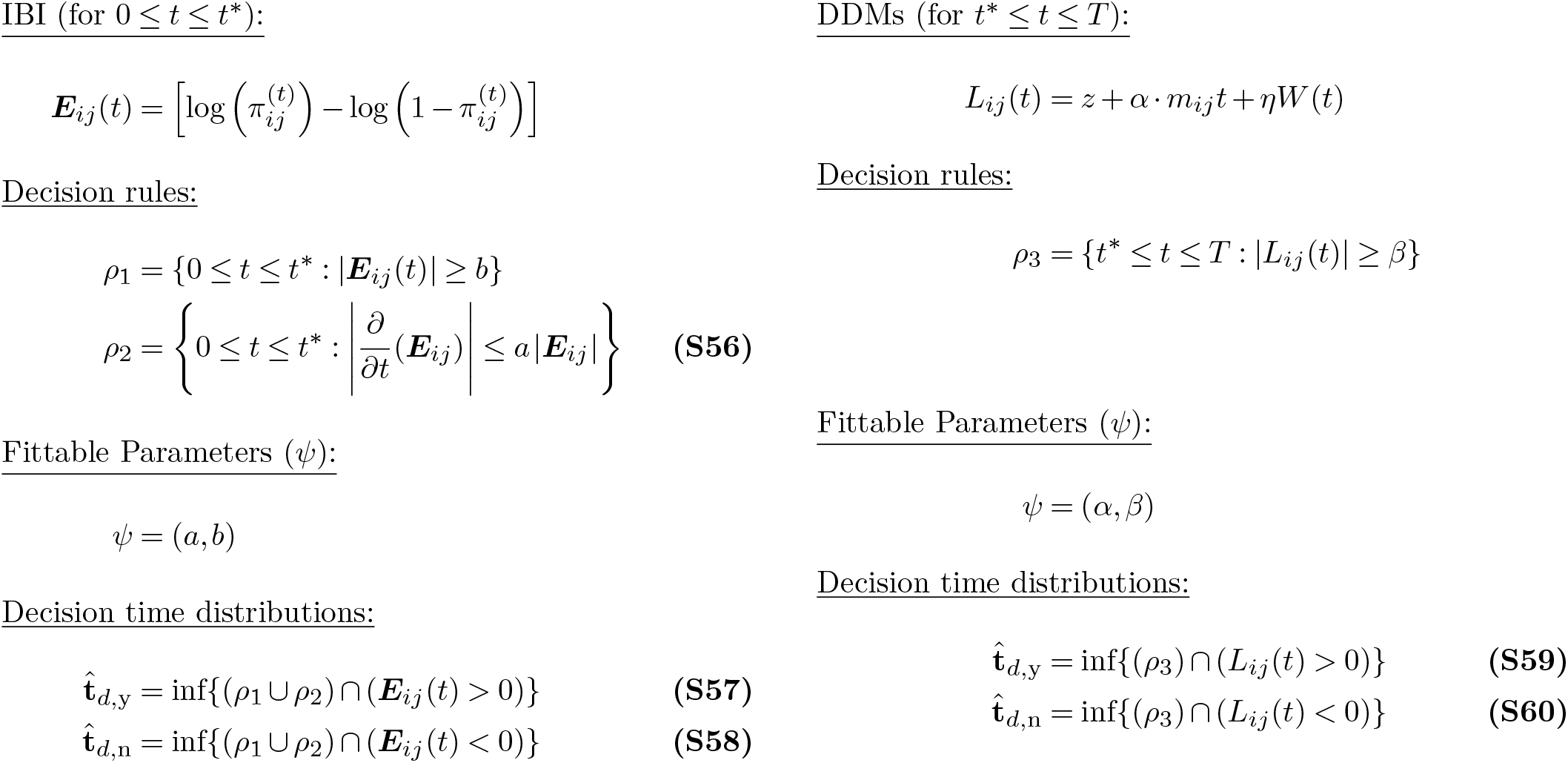
A side-by-side comparison of the iterative and integrative timescales.

Note that neither ***E***_*ij*_(*t*) nor *L*_*ij*_(*t*) are closed-form continuous expressions. However, both the models can be thought of as using a first-order numerical integration procedure to approximate a continuous evidence function.

##### A.3. Types of DDMs

All the DDMs we use can be defined with the following parameterization, where *W*_*ij*_(*t*) indicates that each unique pair (*i, j*) is an independent draw from the Wiener process:

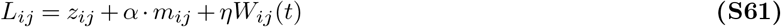

See Table 5 for how we modify substitutions of ε_*ij*_ to create the different DDMs reported in our main text as well as in Supplementary Note 6.

**Table 5.** Types of DDMs. The angle–bracket notation indicates an average. The notation 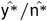 indicates that evidence is averaged per the choice type of the model, at convergence; y: same segment, n: different segments.

| $z_{ij}$ | $m_{ij}$ | Heuristic | Nomenclature in main text |
| --- | --- | --- | --- |
| 0 | 1 | $\alpha$ is decoded from data | Base DDM |
| 0 | $\langle \mathcal{E}_{ij} \rangle_{\hat{y}^*/\hat{n}^*}$ | weight by choice-type | choice-weighted DDM |
| $\mathcal{E}_{ij}$ | $\mathcal{E}_{ij}$ | weight per trial | image-computable DDM/trial-weighted DDM |
| 0 | $\mathcal{E}_{ij}$ | weight drift rate per trial | reduced trial-weighted DDM |
| $\mathcal{E}_{ij}$ | $\langle \mathcal{E}_{ij} \rangle_{\hat{y}^*/\hat{n}^*}$ | weight starting point per trial | reduced trial-weighted DDM |

### Supplementary Note 16: Parameter fitting

In order for the model to generate predictions 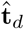 that are most aligned with empirical observations **t**_*d*_, we implemented a parameter fitting procedure. Only two parameters, *ψ* = *a, b* for IBI or *ψ* = *α, β* for DDM, were used for the fitting while the noise *η* remained fixed.

To be able to compare the model’s output of reaction time (which is based on an iteration index) with the continuous reaction time (based on physical time) from humans performing the task, we apply the following transformation to both model and human reaction times: 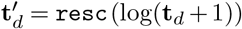. Where the <monospace>resc</monospace>(**x**) function is simply (**x** *−***x**_min_)*/*(**x**_max_*−* **x**_min_).

The goal is to maximize the likelihood that a human decision time **t**^*′d*^ comes from the distribution of model reaction times 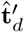. State-of-the-art DDMs use an analytical solution to construct the PDF and compute a loss 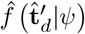 ^35,36^. While this would be possible for the base DDM model, it is not applicable to all models so we opted to empirically construct CDFs for loss instead: 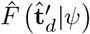. We can use the following empirical likelihood function to calculate the likelihood that a distribution of human reaction times 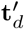 comes from the model CDF 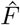^37^:

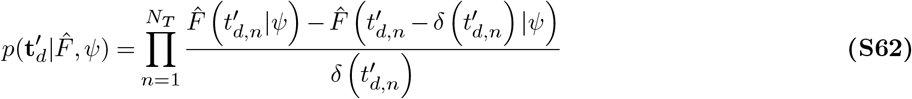

where the product is over all trials, and 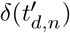 is the time difference with the closest observation smaller than 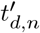. For numerical stability, we optimize the negative log–likelihood 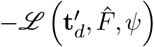.

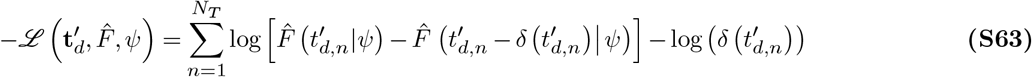

The second term is not a function of the parameters so we ignore it. For the final loss function, we apply the above equation independently to each set of choices

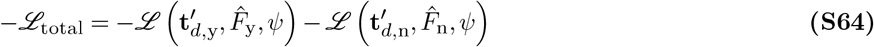

Note that the subscripts y and n indicate the “same segment” subset, and the “different segments” subset as defined by human choices, ignoring the choices of the model. This is because the choices made by the model at specific trials were not expected to match with the human’s, the objective was simply to match reaction time distributions per choice type. equation S64 is not differentiable with respect to the parameters. Therefore we used Markov-Chain Monte-Carlo (MCMC) global minimization methods ^38,39^. Starting from an initial guess (e.g., for IBI, *ψ*^(0)^ = *a*^(0)^, *b*^(0)^) we compute the negative log-likelihood*− ℒ*_total_, which is then iteratively minimized. We heuristically set the following bounds for the MCMC optimizer on parameters in *ψ*: *a∈* (0, 1], *b∈* (0, 5], *α ∈* (0, 50]), *β ∈* (0, 5]. We also found that the minimization of loss improved when multiple initial guesses of *ψ* were used. We used two different initial guesses. By computing this minimization we are finding parameters that maximize the likelihood of human decision times being predicted by our model. For fair comparison across models we cross-validated each model fit with a 5-fold cross validation per case. *− ℒ*_total_ is the loss for one case (*i*.*e*. one participant and one image).

To compare two models (say, A versus B) across cases (one case is one participant and one image), we average their log-likelihood difference ([ℒ_*i,f*_]_A_ − [ℒ_*i*.*f*_]_B_)across cases indexed by *i* and folds indexed by *f*.

### Supplementary Note 17: Movie legend

We provide 3 movie supplements to help with visualization of dynamic segmentation. The image stimulus is the example used in Fig. 1a. The three movies show: 1) the standard IBI algorithm output `[segment_probability_panels.mp4]`(as in Fig. 3b), 2) the no local connectivity ablation output (as in Fig. 7d) `[segment_probability_panels_nlc.mp4]`, and 3) the random initialization output (as in Fig. 7b, middle row) `[segment_probability_panels_rand.mp4]`.

Movie panels: **Top left**: Segment probabilities visualized as in Fig. 7b, coloring is arbitrary, and colors are de-saturated according to their probability. **Top center**: Same as top left, with areas of changing probability marked with contours. Interestingly, the model with local connectivity and perceptually-motivated initialization shows the most stability at late iterations. **Top right**: Contours illustrating where probability is greater than 0.5. Some regions spread outward while others contract. **Bottom left**: Segment probability mass, segment probabilities per segment were summed across all 65 356 pixels. Some segments start out with higher probability mass and decrease towards convergence, while others increase. When initialization is uniformly random, all segments start with the same initial probability mass. **Bottom center**: Segment labels overlaid on the original image, with labels assigned according to the maximum *a posteriori* segment assignment. Comparing to segment probability panels (top left, top center) more structure is visible when visualizing segment probabilities compared to visualizing maximum *a posteriori* segment assignments. **Bottom right**: Segment assignment entropy *H*.

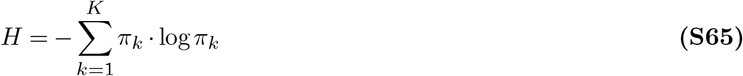

Orange values indicate maximum entropy, purple values indicate minimum entropy. When local connectivity is conserved, entropy is concentrated as perceptual borders. Without local connectivity, entropy is further dispersed.

## Notes

### Competing Interest Statement

The authors have declared no competing interest.

### Summary of Updates

Parameter-free qualitative results from new simulations added; section on ablations of iterative Bayesian inference clarified; supplement updated.

## Bibliography

1. Wagemans, J. et al. A century of gestalt psychology in visual perception: II. conceptual and theoretical foundations. Psychol. Bull. 138, 1218–1252 (2012).

2. Coen-Cagli, R. & Mamassian, P. Are we ready to tackle perceptual segmentation of natural scenes? Vision Res. 240, 108749 (2025).

3. Jolicoeur, P., Ullman, S. & Mackay, M. Curve tracing: a possible basic operation in the perception of spatial relations. Mem. Cognit. 14, 129–140 (1986).

4. Pringle, R. & Egeth, H. E. Mental curve tracing with elementary stimuli. J. Exp. Psychol. Hum. Percept. Perform. 14, 716–728 (1988).

5. Houtkamp, R. & Roelfsema, P. R. Parallel and serial grouping of image elements in visual perception. J. Exp. Psychol. Hum. Percept. Perform. 36, 1443–1459 (2010).

6. Pooresmaeili, A. & Roelfsema, P. R. A growth-cone model for the spread of object-based attention during contour grouping. Curr. Biol. 24, 2869–2877 (2014).

7. Ekman, M., Roelfsema, P. R. & de Lange, F. P. Object selection by automatic spreading of top-down attentional signals in V1. J. Neurosci. 40, 9250–9259 (2020).

8. Vecera, S. P. & Farah, M. J. Does visual attention select objects or locations? J. Exp. Psychol. Gen. 123, 146–160 (1994).

9. Korjoukov, I. et al. The time course of perceptual grouping in natural scenes. Psychol. Sci. 23, 1482–1489 (2012).

10. Jeurissen, D., Self, M. W. & Roelfsema, P. R. Serial grouping of 2d-image regions with object-based attention in humans. Elife 5 (2016).

11. Adeli, H. et al. Human-like object grouping in self-supervised vision transformers. CoRR abs/2603.13994 (2026). URL 10.48550/arXiv.2603.13994.

12. Roelfsema, P. R. Solving the binding problem: Assemblies form when neurons enhance their firing rate-they don’t need to oscillate or synchronize. Neuron 111, 1003–1019 (2023).

13. Goetschalckx, L. et al. Computing a human-like reaction time metric from stable recurrent vision models. Advances in Neural Information Processing Systems 36, 14338–14365 (2023).

14. Marić, M. & Domijan, D. Neural dynamics of spreading attentional labels in mental contour tracing. Neural Netw. 119, 113–138 (2019).

15. Liboni, L. H. B. et al. Image segmentation with traveling waves in an exactly solvable recurrent neural network. Proc. Natl. Acad. Sci. U. S. A. 122, e2321319121 (2025).

16. Chen, M. et al. Incremental integration of global contours through interplay between visual cortical areas. Neuron 82, 682–694 (2014).

17. Klink, P. C., Dagnino, B., Gariel-Mathis, M.-A. & Roelfsema, P. R. Distinct feedforward and feedback effects of microstimulation in visual cortex reveal neural mechanisms of texture segregation. Neuron 95, 209–220 (2017).

18. Franken, T. P. & Reynolds, J. H. Grouping signals in primate visual cortex. Neuron 113, 2508–2520 (2025).

19. Vacher, J., Launay, C., Mamassian, P. & Coen-Cagli, R. Measuring uncertainty in human visual segmentation. PLoS Comput. Biol. 19, e1011483 (2023).

20. Ullman, S. Object recognition and segmentation by a fragment-based hierarchy. Trends Cogn. Sci. 11, 58–64 (2007).

21. Neri, P. Object segmentation controls image reconstruction from natural scenes. PLoS Biol. 15, e1002611 (2017).

22. Bill, J., Gershman, S. J. & Drugowitsch, J. Visual motion perception as online hierarchical inference. Nat. Commun. 13, 7403 (2022).

23. Shivkumar, S., DeAngelis, G. C. & Haefner, R. M. Hierarchical motion perception as causal inference. Nat. Commun. 16, 3868 (2025).

24. Kersten, D., Mamassian, P. & Yuille, A. Object perception as bayesian inference. Annu. Rev. Psychol. 55, 271–304 (2004).

25. Lee, T. S. & Mumford, D. Hierarchical bayesian inference in the visual cortex. J. Opt. Soc. Am. A Opt. Image Sci. Vis. 20, 1434–1448 (2003).

26. Friston, K. A theory of cortical responses. Philos. Trans. R. Soc. Lond. B Biol. Sci. 360, 815–836 (2005).

27. Lengyel, M., Koblinger, Á., Popović, M. & Fiser, J. On the role of time in perceptual decision making. arXiv 1502.03135 (2015).

28. Haefner, R. M., Berkes, P. & Fiser, J. Perceptual decision-making as probabilistic inference by neural sampling. Neuron 90, 649–660 (2016).

29. Drugowitsch, J., Mendonça, A. G., Mainen, Z. F. & Pouget, A. Learning optimal decisions with confidence. Proc. Natl. Acad. Sci. U. S. A. 116, 24872–24880 (2019).

30. Echeveste, R., Aitchison, L., Hennequin, G. & Lengyel, M. Cortical-like dynamics in recurrent circuits optimized for sampling-based probabilistic inference. Nat. Neurosci. 23, 1138–1149 (2020).

31. van Bergen, R. S. & Kriegeskorte, N. Going in circles is the way forward: the role of recurrence in visual inference. Curr. Opin. Neurobiol. 65, 176–193 (2020).

32. Koller, D. & Friedman, N. Probabilistic Graphical Models: Principles and Techniques (MIT Press, 2009).

33. Sun, S., Li, R., Torr, P., Gu, X. & Li, S. Clip as rnn: Segment countless visual concepts without training endeavor. 2024 IEEE/CVF Conference on Computer Vision and Pattern Recognition (CVPR) 13171–13182 (2024).

34. Bogacz, R., Brown, E., Moehlis, J., Holmes, P. & Cohen, J. D. The physics of optimal decision making: a formal analysis of models of performance in two-alternative forced-choice tasks. Psychol. Rev. 113, 700–765 (2006).

35. Gold, J. I. & Shadlen, M. N. The neural basis of decision making. Annu. Rev. Neurosci. 30, 535–574 (2007).

36. Siméoni, O. et al. Dinov3. Trans. Mach. Learn. Res. 2026 (2026). URL https://openreview.net/forum?id=2NlGyqNjns.

37. Dempster, A. P., Laird, N. M. & Rubin, D. B. Maximum likelihood from incomplete data via the em algorithm. J. R. Stat. Soc. Series B Stat. Methodol. 39, 1–22 (1977).

38. Vacher, J., Launay, C. & Coen-Cagli, R. Flexibly regularized mixture models and application to image segmentation. Neural Netw. 149, 107–123 (2022).

39. Neyman, J. & Pearson, E. S. IX. on the problem of the most efficient tests of statistical hypotheses. Philos. Trans. R. Soc. Lond. 231, 289–337 (1933).

40. Wald, A. & Wolfowitz, J. Optimum character of the sequential probability ratio test. Ann. Math. Stat. 19, 326–339 (1948).

41. Fiser, J., Berkes, P., Orbán, G. & Lengyel, M. Statistically optimal perception and learning: from behavior to neural representations. Trends Cogn. Sci. 14, 119–130 (2010).

42. Schwartz, O. & Simoncelli, E. P. Natural signal statistics and sensory gain control. Nat. Neurosci. 4, 819–825 (2001).

43. Coen-Cagli, R., Kohn, A. & Schwartz, O. Flexible gating of contextual influences in natural vision. Nat. Neurosci. 18, 1648–1655 (2015).

44. Farzmahdi, A., Kohn, A. & Coen-Cagli, R. Relating natural image statistics to patterns of response covariability in macaque primary visual cortex. Nat. Commun. 16, 6757 (2025).

45. Hanks, T. D., Mazurek, M. E., Kiani, R., Hopp, E. & Shadlen, M. N. Elapsed decision time affects the weighting of prior probability in a perceptual decision task. J. Neurosci. 31, 6339–6352 (2011).

46. Deneve, S. Making decisions with unknown sensory reliability. Front. Neurosci. 6, 75 (2012).

47. Berkes, P., Orbán, G., Lengyel, M. & Fiser, J. Spontaneous cortical activity reveals hallmarks of an optimal internal model of the environment. Science 331, 83–87 (2011).

48. Girshick, A. R., Landy, M. S. & Simoncelli, E. P. Cardinal rules: visual orientation perception reflects knowledge of environmental statistics. Nat. Neurosci. 14, 926–932 (2011).

49. Stocker, A. A. & Simoncelli, E. P. Noise characteristics and prior expectations in human visual speed perception. Nat. Neurosci. 9, 578–585 (2006).

50. Kar, K., Kubilius, J., Schmidt, K., Issa, E. B. & DiCarlo, J. J. Evidence that recurrent circuits are critical to the ventral stream’s execution of core object recognition behavior. Nat. Neurosci. 22, 974–983 (2019).

51. Martin, D. R., Fowlkes, C. C., Tal, D. & Malik, J. A database of human segmented natural images and its application to evaluating segmentation algorithms and measuring ecological statistics. Proceedings of the Eighth International Conference On Computer Vision 416–425 (2001).

52. Simonyan, K. & Zisserman, A. Very deep convolutional networks for large-scale image recognition. 3rd International Conference on Learning Representations (2015).

53. Cadena, S. A. et al. Deep convolutional models improve predictions of macaque V1 responses to natural images. PLoS Comput. Biol. 15, e1006897 (2019).

54. Cohen, J. A coefficient of agreement for nominal scales. Educational and Psychological Measurement 20, 37–46 (1960). URL 10.1177/001316446002000104.

## References

1. Siméoni, O. et al. Dinov3. Trans. Mach. Learn. Res. 2026 (2026). URL https://openreview.net/forum?id=2NlGyqNjns.

2. Simonyan, K. & Zisserman, A. Very deep convolutional networks for large-scale image recognition. 3rd International Conference on Learning Representations (2015).

3. Cadena, S. A. et al. Deep convolutional models improve predictions of macaque V1 responses to natural images. PLoS Comput. Biol. 15, e1006897 (2019).

4. Schrimpf, M. et al. Integrative benchmarking to advance neurally mechanistic models of human intelligence. Neuron 108, 413–423 (2020).

5. Vacher, J., Launay, C. & Coen-Cagli, R. Flexibly regularized mixture models and application to image segmentation. Neural Netw. 149, 107–123 (2022).

6. Hubert, L. & Arabie, P. Comparing partitions. J. Classif. 2, 193–218 (1985).

7. Wainwright, M. J. & Simoncelli, E. Solla, S., Leen, T. & Müller, K. (eds) Scale Mixtures of Gaussians and the Statistics of Natural Images. (eds Solla, S., Leen, T. & Müller, K.) Advances in Neural Information Processing Systems, Vol. 12 (MIT Press, 1999).

8. Schwartz, O. & Simoncelli, E. P. Natural signal statistics and sensory gain control. Nat. Neurosci. 4, 819–825 (2001).

9. Coen-Cagli, R., Dayan, P. & Schwartz, O. Cortical surround interactions and perceptual salience via natural scene statistics. PLoS Comput. Biol. 8, e1002405 (2012).

10. Coen-Cagli, R. & Mamassian, P. Are we ready to tackle perceptual segmentation of natural scenes? Vision Res. 240, 108749 (2025).

11. Festa, D., Aschner, A., Davila, A., Kohn, A. & Coen-Cagli, R. Neuronal variability reflects probabilistic inference tuned to natural image statistics. Nat. Commun. 12, 3635 (2021).

12. Peel, D. & McLachlan, G. J. Robust mixture modelling using the t distribution. Stat. Comput. 10, 339–348 (2000).

13. Koller, D. & Friedman, N. Probabilistic Graphical Models: Principles and Techniques (MIT Press, 2009).

14. Dempster, A. P., Laird, N. M. & Rubin, D. B. Maximum likelihood from incomplete data via the em algorithm. J. R. Stat. Soc. Series B Stat. Methodol. 39, 1–22 (1977).

15. Bishop, C. M. Pattern Recognition and Machine Learning (Information Science and Statistics) 1 edn (Springer, 2007).

16. Eckford, A. W. The factor graph EM algorithm: applications for LDPC codes. IEEE 6th Workshop on Signal Processing Advances in Wireless Communications, 2005 910–914 (2005).

17. Dauwels, J., Eckford, A. W., Korl, S. & Loeliger, H. Expectation maximization as message passing - part I: principles and gaussian messages. CoRR abs/0910.2832 (2009). URL http://arxiv.org/abs/0910.2832.

18. Lee, T. S. & Mumford, D. Hierarchical bayesian inference in the visual cortex. J. Opt. Soc. Am. A Opt. Image Sci. Vis. 20, 1434–1448 (2003).

19. Beck, J. M., Latham, P. E. & Pouget, A. Marginalization in neural circuits with divisive normalization. J. Neurosci. 31, 15310–15319 (2011).

20. Raju, R. V., Li, Z., Linderman, S. & Pitkow, X. Inferring inference. CoRR abs/2310.03186 (2023). URL 10.48550/arXiv.2310.03186.

21. Neal, R. M. & Hinton, G. E. in A view of the em algorithm that justifies incremental, sparse, and other variants (ed.Jordan, M. I.) Learning in Graphical Models, Vol. 89 of NATO ASI Series 355–368 (Springer Netherlands, 1998).

22. Roweis, S. & Ghahramani, Z. A unifying review of linear gaussian models. Neural Comput. 11, 305–345 (1999).

23. Salakhutdinov, R., Roweis, S. T. & Ghahramani, Z. Optimization with EM and expectation-conjugate-gradient. ICML’03: Proceedings of the Twentieth International Conference on International Conference on Machine Learning 672–679 (2003).

24. Friston, K. A theory of cortical responses. Philos. Trans. R. Soc. Lond. B Biol. Sci. 360, 815–836 (2005).

25. Buxó, C. E. R. & Savin, C. Ranzato, M., Beygelzimer, A., Dauphin, Y. N., Liang, P. & Vaughan, J. W. (eds) A sampling-based circuit for optimal decision making. (eds Ranzato, M., Beygelzimer, A., Dauphin, Y. N., Liang, P. & Vaughan, J. W.) Advances in Neural Information Processing Systems, 14163–14175 (2021).

26. Chen, S., Jiang, L., Rao, R. P. N. & Shea-Brown, E. Oh, A. et al. (eds) Expressive probabilistic sampling in recurrent neural networks. (eds Oh, A.et al.) Advances in Neural Information Processing Systems (2023).

27. Zhang, W.-H., Wu, S., Josić, K. & Doiron, B. Sampling-based bayesian inference in recurrent circuits of stochastic spiking neurons. Nat. Commun. 14, 7074 (2023).

28. Jordan, M. I. & Xu, L. Convergence results for the EM approach to mixtures of experts architectures. Neural Netw. 8, 1409–1431 (1995).

29. Gold, J. I. & Shadlen, M. N. The neural basis of decision making. Annu. Rev. Neurosci. 30, 535–574 (2007).

30. Bogacz, R., Brown, E., Moehlis, J., Holmes, P. & Cohen, J. D. The physics of optimal decision making: a formal analysis of models of performance in two-alternative forced-choice tasks. Psychol. Rev. 113, 700–765 (2006).

31. Fudenberg, D., Newey, W., Strack, P. & Strzalecki, T. Testing the drift-diffusion model. Proc. Natl. Acad. Sci. U. S. A. 117, 33141–33148 (2020).

32. Neyman, J. & Pearson, E. S. IX. on the problem of the most efficient tests of statistical hypotheses. Philos. Trans. R. Soc. Lond. 231, 289–337 (1933).

33. Wald, A. Sequential analysis (John Wiley, 1947).

34. Wald, A. & Wolfowitz, J. Optimum character of the sequential probability ratio test. Ann. Math. Stat. 19, 326–339 (1948).

35. Feller, W. An Introduction to Probability Theory and Its Applications Vol. 1 (Wiley, 1968).

36. Navarro, D. J. & Fuss, I. G. Fast and accurate calculations for first-passage times in wiener diffusion models. J. Math. Psychol. 53, 222–230 (2009).

37. Owen, A. Empirical Likelihood (CRC Press, 2001).

38. Wales, D. J. & Doye, J. P. K. Global optimization by basin-hopping and the lowest energy structures of Lennard-Jones clusters containing up to 110 atoms. J. Phys. Chem. A 101, 5111–5116 (1997).

39. Xiang, Y., Sun, D. Y., Fan, W. & Gong, X. G. Generalized simulated annealing algorithm and its application to the thomson model. Phys. Lett. A 233, 216–220 (1997).

